# Specialization restricts the evolutionary paths available to yeast sugar transporters

**DOI:** 10.1101/2024.07.22.604696

**Authors:** Johnathan G. Crandall, Xiaofan Zhou, Antonis Rokas, Chris Todd Hittinger

## Abstract

Functional innovation at the protein level is a key source of evolutionary novelties. The constraints on functional innovations are likely to be highly specific in different proteins, which are shaped by their unique histories and the extent of global epistasis that arises from their structures and biochemistries. These contextual nuances in the sequence-function relationship have implications both for a basic understanding of the evolutionary process and for engineering proteins with desirable properties. Here, we have investigated the molecular basis of novel function in a model member of an ancient, conserved, and biotechnologically relevant protein family. These Major Facilitator Superfamily sugar porters are a functionally diverse group of proteins that are thought to be highly plastic and evolvable. By dissecting a recent evolutionary innovation in an α-glucoside transporter from the yeast *Saccharomyces eubayanus*, we show that the ability to transport a novel substrate requires high-order interactions between many protein regions and numerous specific residues proximal to the transport channel. To reconcile the functional diversity of this family with the constrained evolution of this model protein, we generated new, state-of-the-art genome annotations for 332 Saccharomycotina yeast species spanning approximately 400 million years of evolution. By integrating phylogenetic and phenotypic analyses across these species, we show that the model yeast α-glucoside transporters likely evolved from a multifunctional ancestor and became subfunctionalized. The accumulation of additive and epistatic substitutions likely entrenched this subfunction, which made the simultaneous acquisition of multiple interacting substitutions the only reasonably accessible path to novelty.

## INTRODUCTION

Many key evolutionary innovations arise from changes to protein sequences that alter their function (Cheng 1998; Zhang et al. 2002; Clark et al. 2003; Dorus et al. 2004; Lunzer et al. 2005; Nielsen et al. 2005; Hoekstra et al. 2006; Christin et al. 2007; Yokoyama et al. 2008; Voordeckers et al. 2012; Projecto-Garcia et al. 2013; Kaltenbach et al. 2018; Jabłońska and Tawfik 2022). Occasionally, these changes stem from dramatic mutational events, including the creation of highly novel coding sequences by gene conversion or ectopic recombination resulting in chimeric proteins (Long and Langley 1993; Nurminsky et al. 1998; Wang et al. 2000; Long et al. 2003; Patthy 2003; Zhang et al. 2004; Ciccarelli et al. 2005; Arguello et al. 2006; Rogers et al. 2010; Rogers and Hartl 2012; Leffler et al. 2017; Méheust et al. 2018; Baker and Hittinger 2019; Brouwers, Gorter de Vries, et al. 2019; Smithers et al. 2019; Baker et al. 2022). While gene conversion can theoretically accelerate the rate of evolution (or even enable adaptation altogether) by bypassing deleterious intermediates, this effect is primarily attributable to the presence of a rugged fitness landscape (Kauffman and Levin 1987; HANSEN et al. 2000; Cui et al. 2002; Bittihn and Tsimring 2017). Such rugged landscapes are manifestations of epistasis in the genotypic combinations underlying the phenotypic map and are prevalent in some empirical systems (Wright 1931; Wright 1932; Maynard Smith 1970; Weinreich et al. 2005; Weinreich et al. 2006; Gong et al. 2013; Weinreich et al. 2013; De Visser and Krug 2014; Sarkisyan et al. 2016; Starr and Thornton 2016; Wu et al. 2016; Pokusaeva et al. 2019; Yi and Dean 2019; Nishikawa et al. 2021; Park et al. 2022; Meger et al. 2024; Metzger et al. 2024). For other proteins, the fitness landscape may be much smoother, meaning that stepwise mutations with additive effects can underlie functional evolution (Lunzer et al. 2005; Bridgham et al. 2006; Weinreich et al. 2006; Poelwijk et al. 2007; Campbell et al. 2016; Kaltenbach et al. 2018; Srikant et al. 2020). In cases where novel protein function is linked to gene conversion events between homologs, these observations therefore raise a fundamental question: are such dramatic mutational events <u>required</u> to evolve new function, or are they probabilistic shortcuts in the evolutionary process whose prevalence is a predictable function of their combined effect size and relative mutation rate? Answering this question has significant implications for understanding and predicting evolutionary trajectories, as well as for designing and engineering novel proteins with desirable functions.

Recently, several remarkably parallel cases of functional innovation have been linked directly or speculatively to gene conversion events in an ecologically and biotechnologically relevant protein family: maltose transporters in *Saccharomyces* yeasts (Baker and Hittinger 2019; Brouwers, Gorter de Vries, et al. 2019; Hatanaka et al. 2022). This protein family consists of transporters similar to the *Saccharomyces cerevisiae* Mal31 protein, which has high specificity and high affinity for the disaccharide maltose, which contains two glucose moieties (Cheng and Michels 1991; Stambuk and Araujo 2001; Salema-Oom et al. 2005; Alves et al. 2008; Brown et al. 2010). Mal31-like proteins are encoded in nearly all genomes of *Saccharomyces* and some closely related species, and they are frequently encoded by multiple paralogs within each genome.

Maltose uptake is also mediated by a second family of proteins, which are related to *S. cerevisiae* Agt1. In contrast to the Mal31-like proteins, Agt1 is a generalist α-glucoside transporter with a broad substrate range, but it has generally lower affinity for those substrates (Han et al. 1995; Stambuk et al. 1999; Stambuk et al. 2000; Alves et al. 2008; Trichez et al. 2019). Notably, Agt1 can transport the glucose trisaccharide maltotriose, a molecule that is biochemically similar to maltose but contains a third glucose moiety. Although sometimes referred to as Mal11, Agt1 is a functionally distinct protein with ≈57% amino acid sequence identity to the Mal31-like proteins. In contrast to the Mal31-like proteins, Agt1-like proteins are rarer, both in presence and in paralog number, in the genomes of *Saccharomyces* yeasts and close relatives (Duval et al. 2010; Horák 2013).

The α-glucoside transporters (Agts) of *Saccharomyces* include the Agt1-like (“generalist”) and Mal31-like (“high-specificity”) proteins, as well as Mph2/3-like proteins (Day et al. 2002), which also have high specificity, albeit for the α-glucoside turanose (Brown et al. 2010). These Agts have been extensively studied due to their important role in the production of beer. Maltose and maltotriose are the two most abundant sugars in brewer’s wort (Meussdorfer and Zarnkow 2009), and their transport into the cell is the rate-limiting step in their fermentation (Zastrow et al. 2001; Horák 2013). The rarity of maltotriose transporters, such as Agt1, which almost always manifests as an inability to ferment this carbon source, therefore presents a barrier to the use of many non-domesticated yeasts in brewing applications.

This barrier is exemplified in *Saccharomyces eubayanus*, the wild, cold-tolerant parent of industrial lager-brewing hybrids (Libkind et al. 2011), whose development for commercial brewing is of great interest (Gibson et al. 2017; Hittinger et al. 2018; Cubillos et al. 2019). As almost all strains of *S. eubayanus* lack generalist Agts capable of transporting maltotriose (Brickwedde et al. 2018; Brouwers, Brickwedde, et al. 2019; Bergin et al. 2022), multiple attempts have been made to evolve maltotriose transporters de novo in *S. eubayanus* strains, using both mutagenesis (Brouwers, Gorter de Vries, et al. 2019) and adaptive laboratory evolution (Baker and Hittinger 2019). These experiments, performed independently in different backgrounds of *S. eubayanus*, yielded results that were as remarkable in their similarity as they were unexpected. In both cases, ectopic gene conversion between paralogous high-specificity (Mal31-like) maltose transporters without any native maltotriose transport capacity (Brickwedde et al. 2018; Baker and Hittinger 2019) resulted in chimeric proteins capable of transporting maltotriose.

Lending weight to the notion that recombination may be a common mechanism by which transporters in the high-specificity Agt family evolve new function, two newly discovered *S. cerevisiae* transporters (Hatanaka et al. 2022), as well as the Mty1 protein (Dietvorst et al. 2005; Salema-Oom et al. 2005), may possess signatures of more ancient gene conversion events (Brouwers, Gorter de Vries, et al. 2019). All these proteins transport maltotriose, but they cluster with Mal31-like proteins in phylogenetic analyses (Baker and Hittinger 2019; Hatanaka et al. 2022). Nonetheless, it remains unclear whether these dramatic mutational events are required for the evolution of novel function in this family or whether they are simply enriched due to the dynamic nature of the subtelomeric regions in which these genes reside (Mefford and Trask 2002; Fairhead and Dujon 2006; Gordon et al. 2009; Brown et al. 2010; Yue et al. 2017; Peter et al. 2018; Liu et al. 2019; O’Donnell et al. 2023).

The yeast α-glucoside transporters are H^+^ symporters belonging to the sugar porter family (TCDB: 2.A.1.1) of the Major Facilitator Superfamily (MFS), a vast, ubiquitous, and ancient group of transmembrane proteins present in all domains of life (Marger and Saier 1993; Pao et al. 1998; Saier 2000; Wang et al. 2020; Saier et al. 2021). Across great evolutionary distances, sugar porters share the highly characteristic MFS fold consisting of twelve transmembrane helices (TMHs) surrounding a hydrophilic central cavity that constitutes the transport channel (Abramson et al. 2003; Guan and Kaback 2006; Sun et al. 2012; Deng et al. 2014; Quistgaard et al. 2016; Bosshart and Fotiadis 2019; Kaback and Guan 2019; Paulsen et al. 2019; Drew et al. 2021). These TMHs are organized into two pseudosymmetrical six-helix bundles (N- and C-terminal), which are separated by a long intracellular linker (ICH domain). The transport channel is surrounded by four helices from each bundle, and TMHs stack tightly against their intra-bundle partners, with additional contacts between the N- and C-terminal domains at the inter-bundle interface. In *S. cerevisiae* Agt1, the sugar substrate and/or proton are thought to be bound primarily by charged residues projecting into this central cavity, which are conserved across fungal Agts (Henderson and Poolman 2017; Trichez et al. 2019). More generally, substrate affinity and specificity in MFS sugar transporters are mediated by extensive hydrogen bonding and occasionally by hydrophobic interactions between the sugar and the protein, as well as steric constraints that limit substrate accommodation; moreover, there is a growing appreciation for the fine-scale and occasionally cryptic contributions to affinity by residues within Van der Waals distance of the substrate (Kasahara et al. 1997; Kasahara and Kasahara 1998; Kasahara and Kasahara 2000; Guan and Kaback 2006; Kasahara et al. 2006; Guan et al. 2007; Kasahara et al. 2007; Kasahara et al. 2009; Kasahara and Kasahara 2010; Kasahara et al. 2011; Sun et al. 2012; Deng et al. 2014; Farwick et al. 2014; Deng et al. 2015; Bosshart and Fotiadis 2019; Kaback and Guan 2019; Drew et al. 2021; Guan and Hariharan 2021).

Nonetheless, the extensive and exquisite biochemical study of MFS sugar transporters has almost exclusively focused on the determinants of native substrate binding and affinity in extant proteins, while questions about how such proteins could evolve the capacity to transport a novel substrate de novo have been largely unaddressed. Understanding evolution-informed design principles in this protein family could enable the engineering of desirable properties in tractable proteins, with significant implications for industrial processes, including the fermentation of cellulosic and hemicellulosic biomass into next-generation biofuels and bioproducts (Ha et al. 2013; Farwick et al. 2014; Young et al. 2014; Turner et al. 2016; Hara et al. 2017; Oh et al. 2017; Casa-Villegas et al. 2018; Kim et al. 2018; Nijland et al. 2018; Nijland and Driessen 2020; Oh and Jin 2020; de Ruijter et al. 2020).

To this end, we aimed to dissect the molecular genetic basis of novel function in the chimeric *S. eubayanus* maltotriose transporter MalT434. *MALT434* arose from an ectopic gene conversion event between genes encoding two paralogous maltose transporters, MalT3 and MalT4, which resulted in the replacement of approximately 230 base pairs of the *MALT4* gene with the homologous portion of *MALT3* (Baker and Hittinger 2019). Both MalT3 and MalT4 are members of the high-specificity maltose transporter family and incapable of transporting maltotriose (Brickwedde et al. 2018; Baker and Hittinger 2019), suggesting that intramolecular epistasis between their protein regions underlies the emergent maltotriose transport by MalT434. The translocated region of *MALT3* encodes TMH 11 and portions of TMHs 10 and 12 (Fig. 1a), and it introduced 11 nonsynonymous mutations to the protein-coding sequence of *MALT4* (Fig. 1b). All three proteins are predicted to have virtually identical structures across their entire folds (pairwise RMSD=0.955Å) and TMHs 10-12 (0.909Å, Fig. S1), suggesting that novel substrate transport might stem from a specific combination of substrate-interacting residues from distal protein regions in MalT434, rather than a global change to protein structure. In the simplest model, as few as a single interacting residue from each protein region could underlie the emergence of novel function, which would make the evolution of new function in this family predictable and tunable; in the most complex model, all 120 amino acid differences between the two parental transporters could contribute, which would render the evolution of new function incredibly difficult.

**Figure 1.**
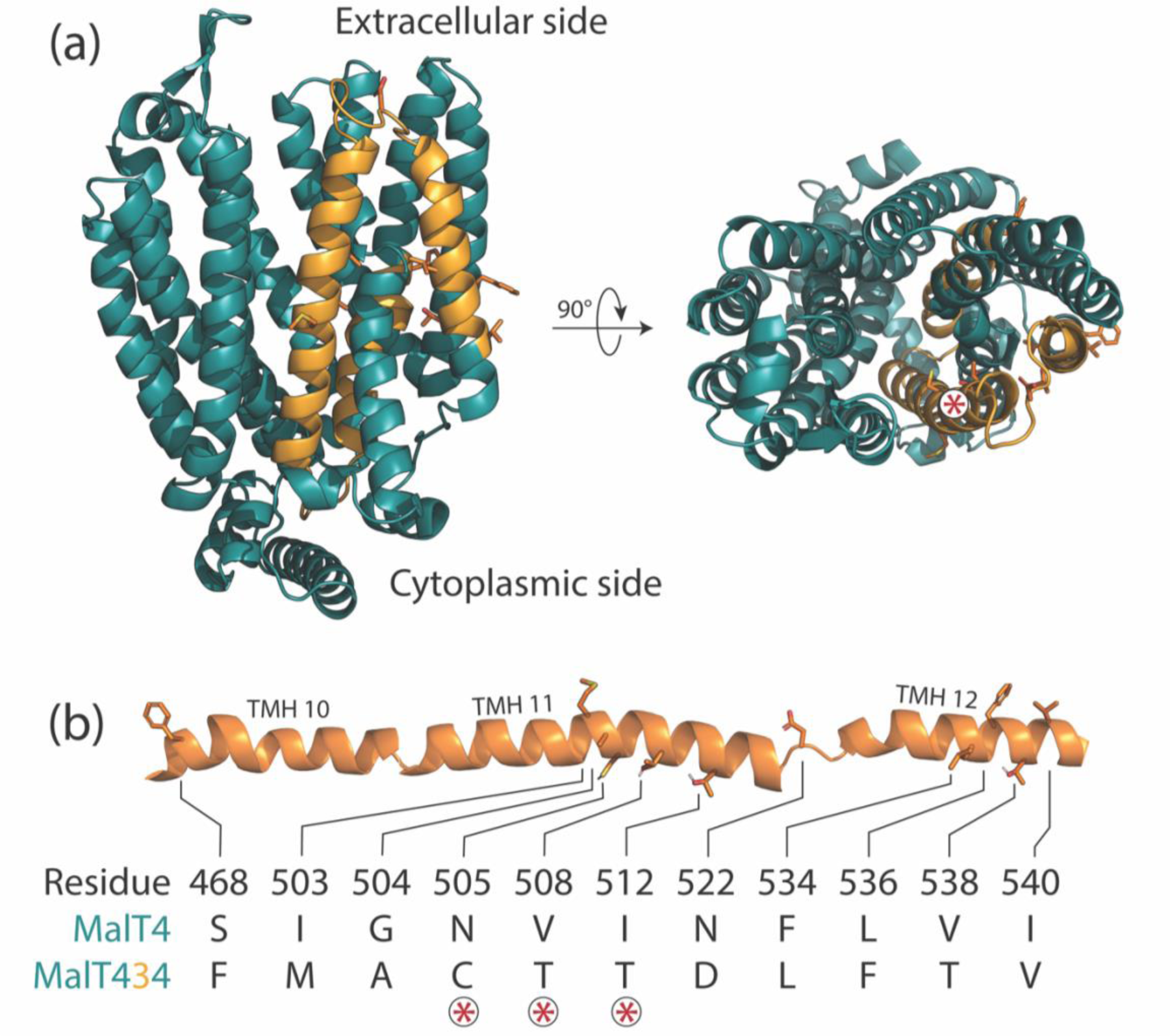
Architecture of a chimeric neofunctionalized α-glucoside transporter. (a) A structural model of the chimeric transporter MalT434 is shown from the side and top views, with alternating colors demarking regions contributed by different parental proteins. The top view is orientated looking down the transport channel. MalT3 side chains are drawn for the 11 substitutions between MalT4 and MalT434. The asterisk label marks the position of the three substitutions on a helical face that bounds the transport channel. (b) Schematic of mutations. The 11 substitutions between MalT4 and MalT434 are drawn as side chains along the cartoon secondary structure of the protein, with loops that connect transmembrane helices truncated for clarity. Polar hydrogens are shown. Asterisks mark the amino acids that face the transport channel.

Here, we show that the basis of maltotriose transport is remarkably complex in this model neofunctionalized transporter. Novel function is shaped by a combination of additive and non-additive interactions between as many as seven regions in the MalT4 backbone and six substitutions across TMHs 10 and 11. At one critical site, very few amino acids can support novel function, which further limits the evolutionary paths available to the wild-type protein; at other sites, these requirements are less stringent. We propose that, overall, novel substrate transport is enabled by widening the transport channel while simultaneously creating a favorable electrostatic environment for the bulkier trisaccharide molecule. Finally, we reconstruct the evolutionary history of the high-specificity and generalist yeast Agts and their relationships to other sugar porters; unexpectedly, we show that the specialist maltose transporters are likely derived and subfunctionalized from a generalist ancestor. This specialization likely involved a gradual refinement of the transport channel to specifically accommodate maltose with higher affinity, which makes the reacquisition of ancestral generalist function difficult to achieve. While our results indicate that rational engineering for novel substrate transport in this protein family is likely to be difficult, they also highlight the abundance and diversity of transporters in biotechnologically relevant yeast species, which could be readily mined for desirable functions that have been exquisitely refined over billions of years of evolution, as well as perhaps recombined into new functions.

## RESULTS

### High-order intramolecular interactions are required to evolve a novel function in maltose transporters

We first investigated the scope and complexity of intramolecular interactions shaping the emergence of novel function in MalT434. We coarsely defined functional protein regions as the twelve transmembrane helices (TMHs), the intracellular (ICH) domain, and the partially unstructured intracellular N- and C-terminal regions. We iteratively constructed novel chimeric genes encoding transporters from MalT3 and MalT4 components and tested their ability to support growth on maltotriose when expressed from the native *MALT4* locus (Fig 2). Unsurprisingly, the C-terminal portion of MalT4 present in MalT434 was neither necessary (construct 1) nor sufficient (construct 17) for maltotriose transport; indeed, its replacement with the corresponding region of MalT3 improved growth on maltotriose by 15.3% (*p* = 5.3×10^−4^, Mann-Whitney *U* test). By contrast, replacement of TMHs 8 and 9 and the N-terminal half of TMH 10 with their MalT3 counterparts (construct 2) reduced growth by 11.6% compared to MalT434 (*p* = 0.184), while still supporting robust growth. Dissection of the region N-terminal to TMH 8 revealed that the key interaction enabling maltotriose transport occurs between TMHs 10 and 11 of MalT3 and TMH 7 from MalT4. While necessary, this region alone was not sufficient to enable maltotriose transport in every protein context. In addition to the epistatic interaction between TMHs 7, 10, and 11, growth on maltotriose required the presence of TMHs 1 and 2 from MalT4 in combination with the ICH domain from MalT3 (construct 7), or alternatively, one or more of TMH 5, TMH 6, and the ICH domain from MalT4 (construct 15).

**Figure 2.**
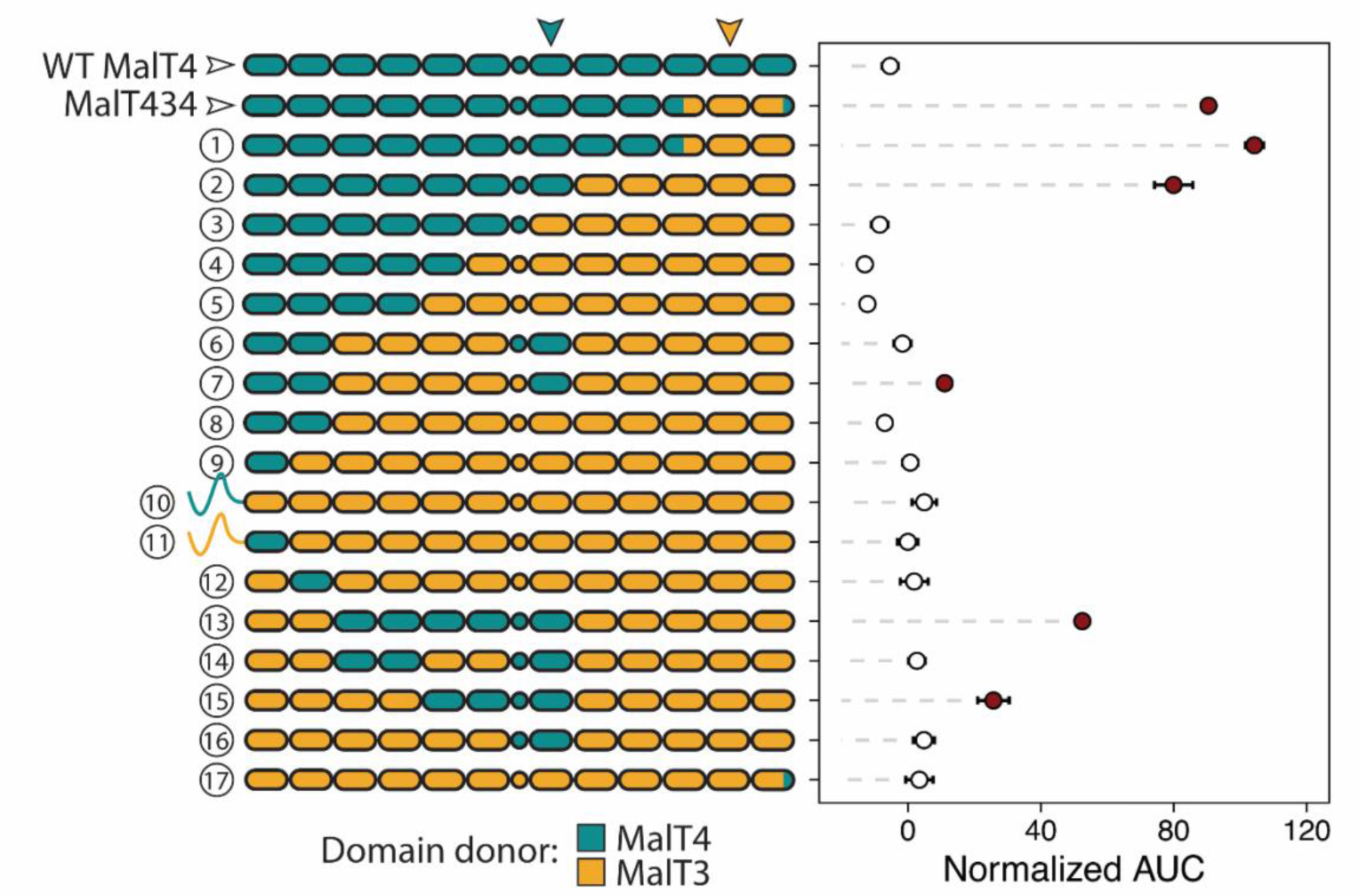
High-order intramolecular interactions are required to evolve a novel function in chimeric α-glucoside transporters. Points and bars show mean +/− SEM of normalized growth on maltotriose (AUC, area under the curve) of strains expressing chimeric transporters or wild-type MalT4 (top row). Filled circles denote growth significantly greater than the negative control (*p* < 0.01, Mann-Whitney *U* test with Benjamini-Hochberg correction). The architecture of each tested transporter is depicted as a cartoon on the y-axis, where rounded rectangles represent each of the twelve transmembrane helices and circles represent the intracellular ICH domain that links the N- and C-terminal six-helix bundles; regions are colored by parental protein identity. In almost every case, the N- and C-terminal intracellular regions have the same parental protein identity as the neighboring transmembrane helix and are omitted for clarity; the two exceptions are depicted. Inverted arrows indicate the location and identity of protein regions underlying the largest detected intramolecular interaction.

For chimeric constructs containing potentiating sequences at TMHs 5-7 and 10-12, growth on maltotriose generally increased additively with the number of MalT4 regions incorporated (linear regression, *p* < 2.2×10^−16^). Nonetheless, we found significant support (ANOVA, *p* < 2.2×10^−16^) for pairwise epistasis between the tested protein regions, including in the sign of the effects of the ICH domain and the C-terminal region (residues 541-613). For example, the addition of TMH 3 and TMH 4 from MalT4 in conjunction with MalT4 TMH 7 only increased growth on maltotriose if TMH 5 and TMH 6 from MalT4 were also present; similarly, the addition of TMH 1, TMH 2, and the ICH domain from MalT4 in conjunction with TMH 7 did not improve growth (construct 6 vs. 16, Fig. 2) unless in the presence of TMHs 3-6 from MalT4 (construct 2 vs. 13, 52% increase, *p* = 2.4×10^−4^). Along the quantitative functional spectrum of MalT3/4 chimeric proteins enabling growth on maltotriose, we therefore detected a complex combination of additive and epistatic intramolecular interactions among at least six protein regions.

### Numerous substitutions are required to evolve a novel function in maltose transporters

We next dissected the contributions of the 11 substitutions in MalT434 relative to MalT4 (Fig. 1b) by introducing subsets of these to the gene encoding the native MalT4 protein (Fig. 3). We first tested the effect of a pair of suggestive substitutions, S468F and N522D, which were both unique in their location in the 3D structure and differed notably in side-chain chemistry. Nonetheless, this pair of mutations was insufficient for novel function in MalT4, so we coarsely tested the effect of the sets of mutations occurring before and after the end of TMH 11. Introduction of the five substitutions from residues 522-540, which span an extracellular loop and the majority of TMH 12, was insufficient to confer any growth on maltotriose. By contrast, the six mutations affecting TMHs 10 and 11 were sufficient to confer growth on maltotriose, and even improved it by 13.3% relative to MalT434 (*p* = 5.6×10^−7^, Mann-Whitney *U* test). Within this contiguous patch of substitutions, however, the contribution of individual amino acids to novel function was remarkably complex. Reversion of the six mutations singly to their MalT4 identity revealed that each had a significant effect on maltotriose growth, ranging from a 23.5% reduction (A504G, *p* = 2×10^−6^) to its complete abrogation (C505N, *p* = 5.2×10^−11^), with an average effect of 57.1%. We detected significant (*p* < 2.2×10^−16^) evidence of pairwise epistasis between substitutions, regardless of whether we considered all 11 sites or only the 6 on TMHs 10 and 11. Epistatic effects were notably non-uniform among tested combinations: for example, two single reversion mutations (M503I and T508V) had similar effects of 49.1% (*p* = 3.2×10^−7^) and 44.1% (*p* = 9.1×10^−13^) when introduced in the six-substitution background that supported robust growth on maltotriose. By contrast, when introduced in a four-mutation background with reduced ability to support growth on maltotriose (M503 C505 T508 T512), the effect of M503I remained large (42.6%, *p* = 0.002), while T508V effected only a small further reduction (4.97%, *p* = 0.8). Overall, we found that establishing novel function in MalT4 required a combination of three amino acid substitutions only accessible through a minimum of four non-consecutive nucleotide substitutions to the wild-type gene: N505C (2 nucleotide substitutions), I512T (1 substitution), and one of I503M (1 substitution) or V508T (2 substitutions).

**Figure 3.**
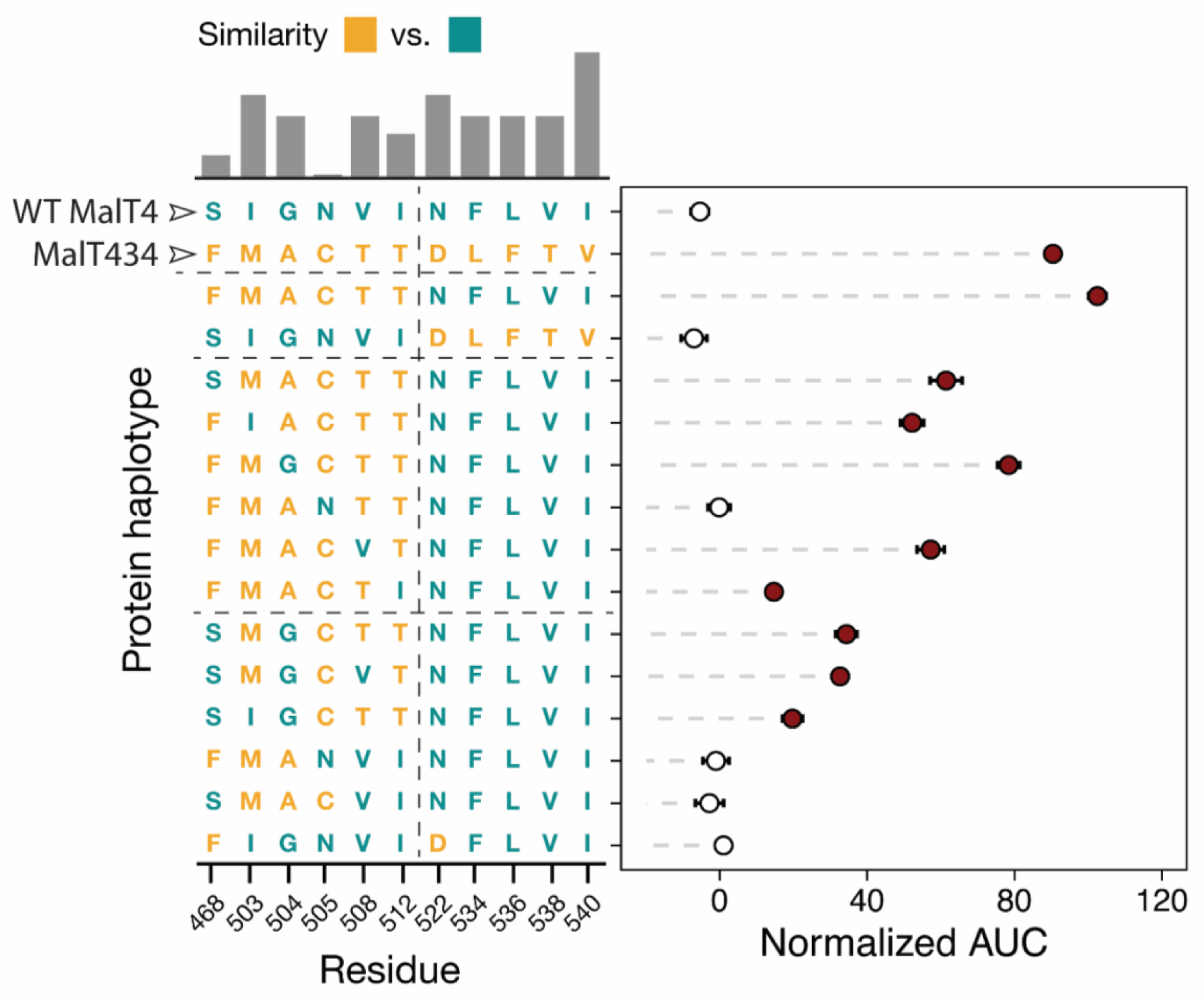
Numerous substitutions are required to evolve a novel function in a maltose transporter. Points and bars show mean +/− SEM of normalized growth on maltotriose (AUC, area under the curve) of strains expressing MalT4 variants. The genotype of each protein at the 11 sites that differ between MalT4 (top row) and MalT434 (second from top row) is depicted on the Y-axis. Filled circles denote growth significantly greater than the negative control (*p* < 0.01, Mann-Whitney *U* test with Benjamini-Hochberg correction). The bar chart shows rescaled BLOSUM similarity between the MalT4 and MalT3 residue at that site, with a higher bar indicating a more conservative substitution. Horizontal dotted lines in the protein haplotype grid separate related groups of genotypes. The vertical dotted line demarcates the substitutions that are sufficient (left) to impart novel function to MalT4 and those that are insufficient (right).

### Granular mapping of epistasis between distal protein regions

Given the size of interacting protein regions and the complexity of their contributions to novel function, we sought to identify the key difference in amino acid sequence responsible for the large epistatic effect of transmembrane helix 7. The two parental transporters differ at six sites along TMH 7 (Fig. S2a): two neighboring substitutions (K357C and V358I, expressed relative to MalT4) occur at the intracellular C-terminal end, while two (A371I, V375T) are located approximately halfway along the helix and likely to be embedded in the plasma membrane. Two (A378T, S379Q) project into or neighbor the transport channel, differ in size and/or polarity, and are in close three-dimensional proximity to mutated residues on TMH 11 in MalT434 (Fig. 4a, Fig. S2b). We reasoned that one or both of A378T and S379Q might have a large effect on the interaction between TMH 7 and the translocated region of MalT3 present in functional chimeric transporters. To test these hypotheses, we mutated each of these residues to their MalT3 identity, singly and in combination, in a gene encoding the MalT4 transporter harboring the six mutations on TMHs 10 and 11 that conferred maximal maltotriose transport (Fig. 4b). While the A378T mutation did not affect growth on maltotriose, S379Q abolished it completely. The large epistatic interaction between TMH 7 and TMH 11 can thus be attributed to a single amino acid.

**Figure 4.**
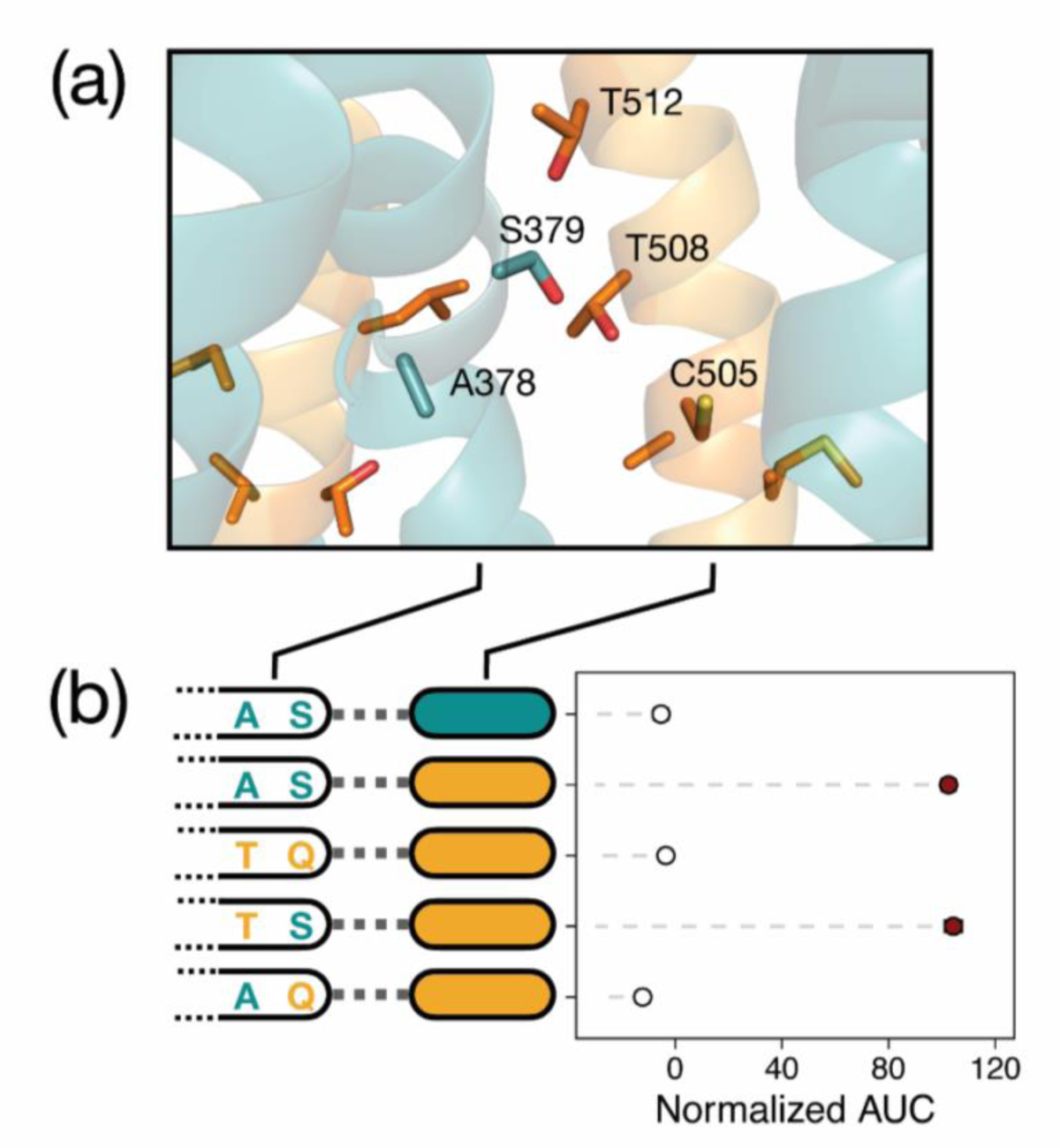
A single amino acid underlies a large epistatic effect. (a) Structural model of MalT434 with helices colored as in Fig. 1. Side chains are drawn for amino acids on transmembrane helices 7, 11, and 12 that are polymorphic between MalT3 and MalT4, and those that are proximal to or project into the transport channel are labeled. (b) Points and bars show mean +/− SEM of normalized growth on maltotriose (AUC, area under the curve) of strains expressing transporter variants. Filled circles denote growth significantly greater than the negative control (*p* < 0.01, Mann-Whitney *U* test with Benjamini-Hochberg correction). For each transporter, the parental protein identity at transmembrane helix 11 (filled rectangular ovals) and residues 378 and 379 in transmembrane helix 7 is depicted.

### Novel transporter function is constrained by specific biochemical requirements and context dependence

The mutational event that generated MalT434, as well as our experiments dissecting it, only sampled variation between two binary states: the specific amino acid residues of the parental proteins at each homologous site. In native contexts, however, many more amino acid substitutions are accessible in mutational space through single- or multi-nucleotide mutations; for example, seven amino acid substitutions require only a single nucleotide change from an asparagine codon, which is the wild-type amino acid at the crucial 505 site. While we found complex interactions between many sites to contribute to novel function in MalT4, the evolution of maltotriose transport would be far less constrained and more accessible through sequential point mutations if biochemically similar amino acids at key sites could enable a degree of novel function because it would increase the mutational target size and pool of mutations conferring a fitness benefit (Miyazaki and Arnold 1999; Podgornaia and Laub 2015).

We thus sought to clarify the biochemical requirements for maltotriose transport in a specific potentiated context: a MalT4 transporter harboring S379, F468, M503, A504, T508, and T512. In this state, amino acid identity at position 505 is crucial with the wild-type asparagine incapable of supporting growth on maltotriose and the recombinant cysteine supporting robust growth (Fig. 3). We successfully mutated this residue to 17 of the 20 possible amino acids, measured their ability to support growth on maltotriose, and used regression analyses to estimate the effect of side chain physicochemical properties on measured function (Fig. 5). Remarkably, only three substitutions supported any degree of statistically significant growth above baseline: serine, glycine, and cysteine. Side chain aromaticity, volume, composition, and hydropathy were all significant (*p* << 0.01) predictors of function, as was overall similarity to the wild-type residue asparagine. Even so, the strengths of these associations were almost entirely driven by the C505 variant: when these data were omitted, the global explanatory power was reduced dramatically (adjusted R^2^: 0.2263 vs. 0.8664; F-statistic: 9.533 vs. 242). Although some physicochemical properties remained statistically significant predictors of function, the strengths of these associations were generally weak (maximum |Kendall’s Τ|: 0.212).

**Figure 5.**
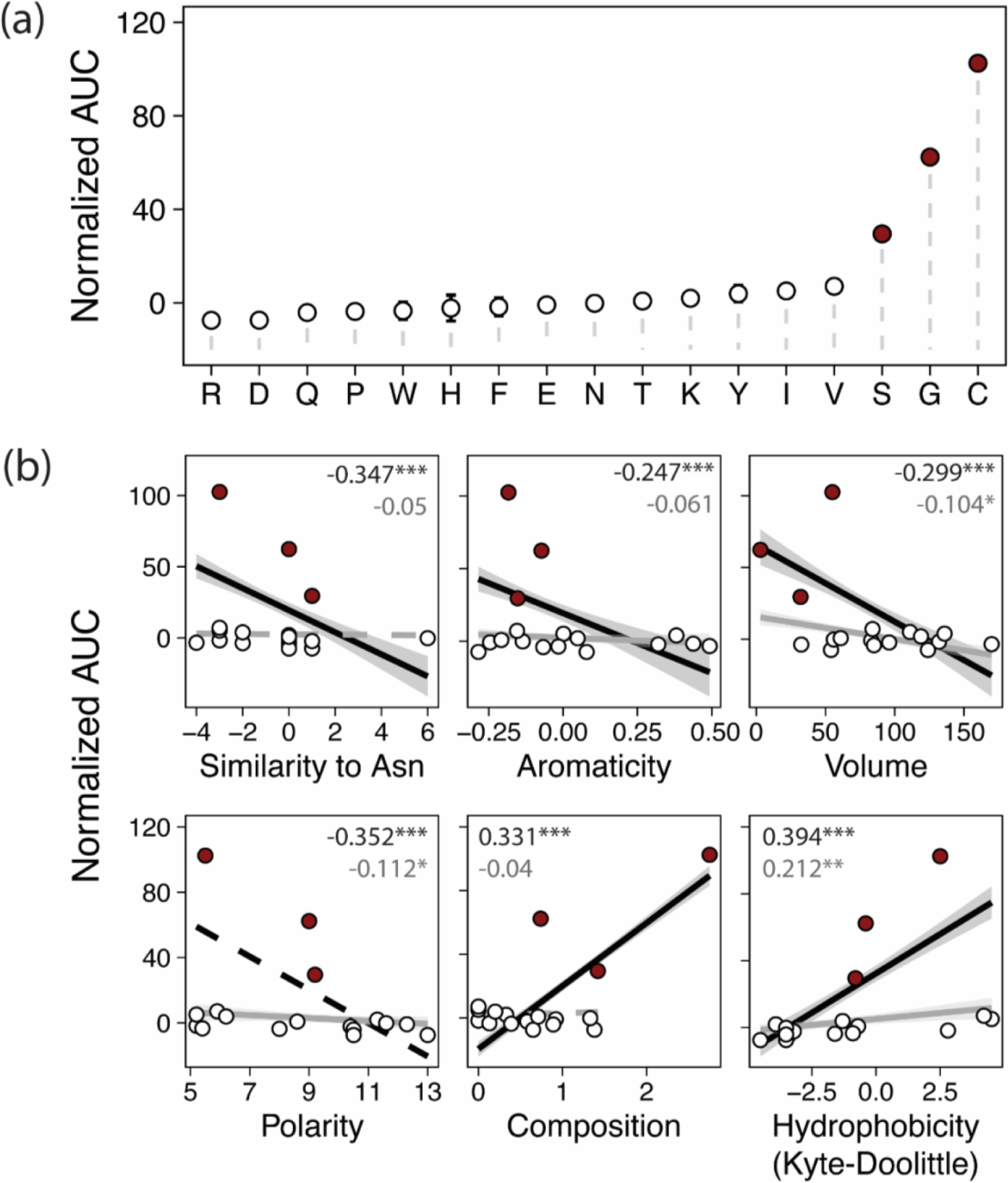
Physicochemical requirements constrain the evolution of novel function. (a) Points and bars show mean +/− SEM of normalized growth on maltotriose (AUC, area under the curve) of strains expressing MalT4 variants. The x-axis shows the amino acid identity at position 505; all variants share F468, M503, A504, T508, and T512. Filled circles denote growth significantly greater than the negative control (*p*<0.01, Mann-Whitney *U* test with Benjamini-Hochberg correction). (b) Correlations between growth and properties of the amino acid variant at position 505. Growth is plotted as in (a) against physicochemical property or overall similarity to the wild-type residue at position 505, asparagine. Lines and shaded ranges show regressions and 95% confidence intervals for significant (*p* < 0.05) regressions for all data (black) or after removing observations for C505 (gray). Dotted lines show regressions that are not statistically significant. Inset text shows Kendall’s Τ; \*\*\**p* < 10^−6^, \*\**p* < 10^−4^, \**p* < 0.05.

Qualitatively, the fine-scale stringency of physicochemical requirements at position 505 was also noteworthy. Glycine, serine, and cysteine are three of the smallest amino acids, but amino acids with similar side chain volumes did not support growth on maltotriose. Serine and cysteine have side chains of similar size and structure capable of forming hydrogen bonds, but they differ in their polarity and hydrophobicity; nonetheless, residues similar to cysteine in both of these metrics did not support novel function. Indeed, C505’s ability to support novel function appeared to be the result of the specific combination of cysteine’s physicochemical properties (Fig. S3), albeit not due to its unique capacity to form disulfide bridges (Drew et al. 2021). Remarkably, this effect was dependent on positional context within the transporter: while substituting cysteine to serine at 505 reduced growth by 71.2% (*p* = 8.8×10^−5^), making the orthogonal serine to cysteine substitution at another key site, S379 (Fig. 4) reduced growth by 17.7% (*p* = 1.9×10^−6^) while still supporting robust growth (Fig. S4). Thus, while serine was largely unable to recapitulate the effect of cysteine at 505, the similarity between the two was sufficient to satisfy the requirements for novel function at position 379. The same was not true of two other hydrogen bond-competent residues, glutamic acid and glutamine, whose introduction at position 379 abolished growth (Fig. S4). This result suggests that, while serine and cysteine are interchangeable at this site, interactions between physical and chemical side chain properties still play a role. Finally, we found further evidence for these fine-scale requirements at position 512, where mutation of the permissive threonine to valine reduced growth by 34.5% (*p* = 7.4×10^−9^), while still supporting significantly improved growth over the wild-type MalT4 residue isoleucine (78.1% increase, *p* = 1.2×10^−6^). In summary, we find that the strengths, stringencies, and bases of physicochemical requirements all vary between sites that are critical for establishing novel function in MalT434. These results suggest that the serendipitous acquisition of a set of epistatically sufficient residues is highly improbable by point mutations alone (Lynch 2005).

### High-specificity transporters are evolutionarily derived

The sum of our molecular analyses suggested that the acquisition of novel substrate transport by the high-specificity maltose transporter MalT4 is highly improbably and accessible only through the simultaneous acquisition of numerous interacting substitutions. This observation is consistent with previous failed attempts to establish a maltotriose transporter by introducing as many as 14 rational mutations to *S. cerevisiae* Mal61 (Hatanaka et al. 2022), a prototypical high-specificity maltose transporter closely related to MalT4. However, the presence of closely related generalist α-glucoside transporters, as typified by *S. cerevisiae* Agt1, suggests that this ability evolved at least once among yeast α-glucoside transporters. We sought to clarify the timing and mode of this historical evolutionary innovation by examining the phylogenetic relationships between the generalist and specialist α-glucoside transporters within Saccharomycotina yeasts, which have previously been assessed on only a few taxa (Brown et al. 2010; Cousseau et al. 2013; Baker and Hittinger 2019; de Ruijter et al. 2020; Hatanaka et al. 2022; Donzella et al. 2023).

We first generated high-quality protein-coding gene annotations for published genomes from 332 yeast species from the model subphylum Saccharomycotina, which spans more than 400 million years of evolution (X.-X. Shen et al. 2018). To formally test the expected monophyly of the α-glucoside transporters within the broader sugar porter family, we retrieved homologs of *S. cerevisiae* sugar porters from these predicted proteomes and constructed a comprehensive phylogeny of these 8,403 ecologically and biotechnologically relevant MFS proteins. This phylogeny split into several major clades, many of which contained at least one functionally characterized protein from *S. cerevisiae* or another species (Fig. S5). Both the high-specificity (Mal31- and Mph2/3-like) and generalist (Agt1-like) α-glucoside transporters clustered in a monophyletic group (“Agt clade”) that excluded other sugar porter families. All proteins in the Agt clade from the newly circumscribed order Saccharomycetales (Groenewald et al. 2023) grouped together with strong support (Fig. 6a). The monophyly of the Saccharomycetales Agts was interrupted in two cases: 1) a single protein from *Ogataea naganishii* sister to the *Lachancea* Agt1-like proteins; 2) and, more notably, a well-supported clade of Agts from *Brettanomyces anomalus* and *Brettanomyces bruxellensis*. The *Brettanomyces* species are documented recipients of numerous horizontal gene transfer events, including for genes involved in the metabolism of sucrose, an Agt1 substrate (Stambuk et al. 2000; Woolfit et al. 2007; Roach and Borneman 2020). Notably, *B. bruxellensis* is commonly associated with brewing environments, where its propensity to vigorously consume diverse sugars and independent evolution of aerobic fermentation make it a frequent contaminant and occasional desired contributor (Rozpedowska et al. 2011; Serra Colomer et al. 2019; Colomer et al. 2020).

**Figure 6.**
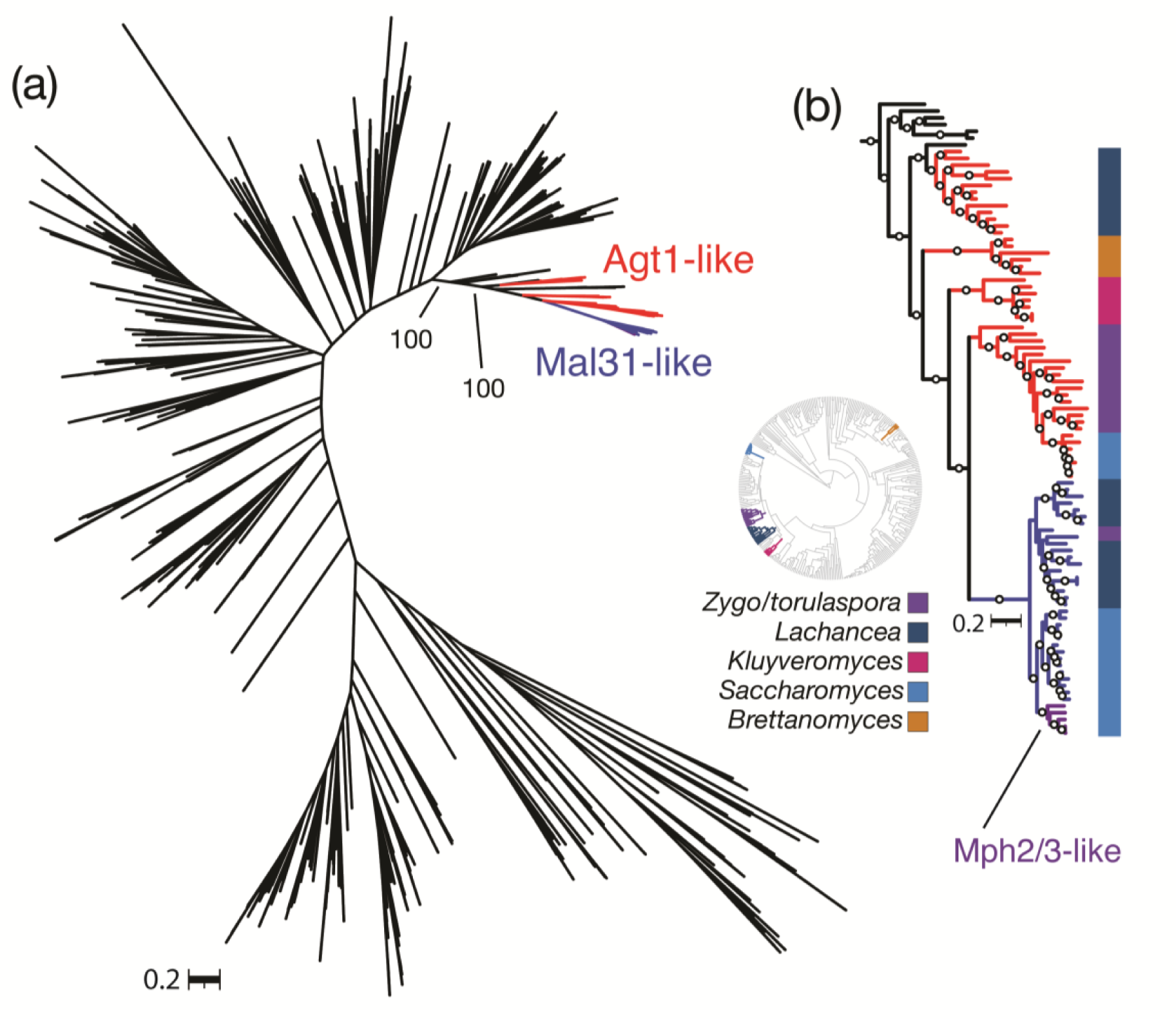
The high-specificity maltose transporters are evolutionarily derived and restricted to a subset of Saccharomycetales. (a) Consensus phylogeny of the α-glucoside transporter clade from 332 budding yeast genomes. Agt1-like and Mal31-like proteins from all Saccharomycetales are colored, as is the *Saccharomyces*-specific Mph2/3 clade. Bootstrap support is shown for two splits leading to the Saccharomycetales. (b) Rooted consensus tree of the clade containing Saccharomycetales α-glucoside transporters. Branches are colored as in (a) with the inclusion of a well-supported clade of *Brettanomyces* Agt1-like proteins that nests within the Saccharomycetales; the *Saccharomyces-*specific Mph2/3 clade is indicated. Circles denote branches with >90% bootstrap support. Colored bars outside the tree show genus-level taxonomic assignment, and the inset circular tree shows the Saccharomycotina species phylogeny (X.-X. Shen et al. 2018) with those genera colored; *Zygo/torulaspora* represents *Zygosaccharomyces, Zygotorulaspora,* and *Torulaspora*. The rooted maximum-likelihood tree can be found in Fig. S6. Newick-formatted trees are available in Data S2 and S3.

Surprisingly, the clade containing high-specificity *Saccharomyces* maltose transporters only included taxa from closely related species in the genera *Saccharomyces* and *Lachancea*, as well as one protein each from *Zygotorulaspora florentina* and *Zygosaccharomyces kombuchaensis* (Fig. 6b). Among the high-specificity Agts, the Mph2/3 clade was further restricted to *Saccharomyces kudriavzevii*, *Saccharomyces mikatae*, *Saccharomyces paradoxus*, and *S. cerevisiae* (Fig. 6b), which is consistent with an origin in the common ancestor of these species following their split from *Saccharomyces arboricola* and a recent segmental duplication in *S. cerevisiae* (*Saccharomyces jurei* is absent in this dataset). The sister clade to the high-specificity proteins contained generalist Agts from *Saccharomyces, Torulaspora*, and *Zygotorulaspora* species, with deeper branches to *Kluyveromyces* and *Lachancea* homologs (Fig. 6b). We thus conclude that the high-specificity transporters typified by *S. cerevisiae* Mal31, including *S. eubayanus* MalT4 and MalT3, form a clade restricted to Saccharomycetales.

### Generalist-like transporters are quantitatively correlated with growth on α-glucosides

Our phylogenetic analyses suggested that the high-specificity Agts are evolutionarily and functionally derived from a generalist ancestor. In this model, the vast array of uncharacterized Agt-clade proteins encoded by diverse yeast species should include generalist transporters or transporters that became subfunctionalized following duplication of a generalist ancestor, and their presence should support growth on substrates of the generalist Agts. We collected quantitative growth measurements for 287 of the 332 species in our phylogenetic dataset on three sugars that are substrates of the generalist transporter *S. cerevisiae* Agt1 but not of the high-specificity transporters: maltotriose, trehalose, and methyl-α-glucoside (Han et al. 1995; Stambuk et al. 1999; Stambuk and Araujo 2001; Alves et al. 2008; Brown et al. 2010). We found many species across the Saccharomycotina to be capable of vigorous growth on these sugars as a sole carbon source (Fig. 7a). Growth on all three α-glucosides was nearly ubiquitous among Serinales, a speciose order with a high incidence of carbon niche-breadth generalists (Opulente et al. 2024). Most notably, growth on maltotriose was widespread across the yeast subphylum, in contrast to the documented rarity of this trait in the model genus *Saccharomyces* (Duval et al. 2010; Gallone et al. 2018; Langdon et al. 2020; Hutzler et al. 2021; Gyurchev et al. 2022; Peris et al. 2023). This metabolic deficiency was concomitant with the paucity of generalist-like Agt proteins encoded in Saccharomycetales genomes, which was similarly not representative of other yeast orders (Fig. 7b; *p* = 1.9×10^−13^). Indeed, patterns of α-glucoside growth qualitatively tracked the presence of genes encoding Agt proteins, with both subject to clear evolutionary shifts including losses (e.g. Saccharomycodales, Sporopachydermiales, and Trigonopsidales; *Saturnispora*, *Zygosaccharomyces*, *Eremothecium*, *Kazachstania, Nakaseomyces*, *Naumovozyma*, and *Tetrapisispora* spp.) and amplifications (*Debayromyces*, *Metschnikowia,* and *Kuraishia* spp.; subclades of Phaffomycetales, Dipodascales, Pichiales, and Lipomycetales). We used phylogenetically corrected least squares regressions (PGLS) to statistically test the strength of the correlation between Agt count and growth on each of the three tested Agt1 substrates (Fig. 7c). We detected significant positive correlations between Agt count and growth on each of the three α-glucosides (*p* ≤ 0.007). Thus, the generalist-like Agts detected in most Saccharomycotina genomes are likely to be true generalist transporters or recently subfunctionalized derivatives.

**Fig. 7.**
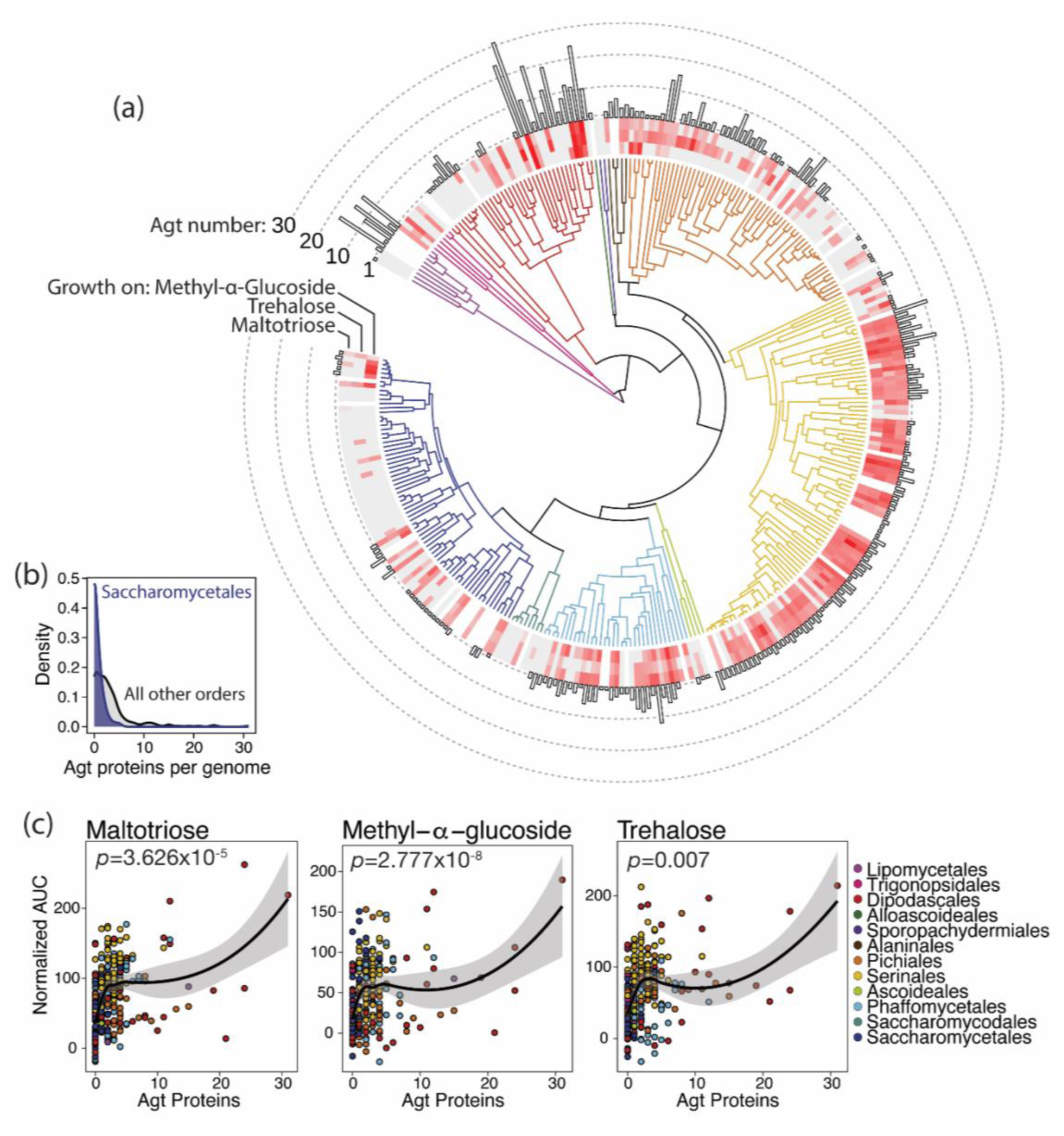
Species with Agt proteins grow on Agt1-specific substrates. (a) Time-calibrated phylogeny of 332 Saccharomycotina species (X.-X. Shen et al. 2018) with branches colored (key in panel c) by taxonomic order (Groenewald et al. 2023). Heatmaps around the tree show growth (normalized area under the curve) on α-glucosides: methyl-α-glucoside (inner ring), trehalose (middle ring), and maltotriose (outer ring). Gray boxes denote no growth above background; white boxes represent unsampled species. The bar chart shows the number of proteins in the α-glucoside transporter clade for each genome. (b) Generalist Agt content of Saccharomycetales genomes is not representative. Density plots show distributions of the number of Agt-clade proteins per genome for Saccharomycetales species (blue density) and species from all other orders (gray). (c) Scatterplots of Agt-clade transporter count versus growth on each α-glucoside. Each species is represented by a point, colored by taxonomic order. Lines and shaded regions are loess-smoothed regressions of the untransformed data; inset *p*-values are from phylogenetically corrected regressions (PGLS).

## DISCUSSION

In the present work, we sought to understand how novel function could evolve in a model yeast α-glucoside transporter. To this end, we dissected the molecular basis of maltotriose transport in MalT434, which represents one of the most evolutionarily recent functional innovations in this family. We found that, in this chimeric protein, novel function is an emergent property of extensive additive and non-additive interactions between multiple protein regions and multiple residues on TMHs 7, 10, and 11 (Figs. 2-4). We observed that even conservative amino acid changes, as well as residues not predicted to interact with the substrate, had significant and unexpected effects on maltotriose transport (Fig. 3, Fig. 5). We also found evidence that the stringency of side chain physicochemical requirements likely differs substantially between crucial residues (Fig. 5, Fig. S4). Taken together, these results demonstrate that the evolution of novel function in a high-specificity Agt is highly constrained, which is consistent with recent observations (Hatanaka et al. 2022). In this model, the evolution of novel function in this family by gene conversion may indeed be the only remotely probable way that all the necessary interacting residues can readily be assembled in a single molecule, even if paralogs are free to sample neutral or deleterious mutational steps.

The gene conversion events leading to novel function in high-specificity yeast Agts share striking parallelism at both the sequence and structural scales. For example, the portions of Mty1 inferred to derive from different parental proteins encompass many of the same regions that we identified as having crucial interactions in MalT434 (Fig. S7a). Even more strikingly, the homologous residues at five of the seven sites that affect maltotriose transport in MalT434 are conserved in Mty1 (Fig. S7b). At the other two sites, Mty1 possesses amino acids that support reduced, but significant, growth in MalT434 (C505S and T512I). While many of the same sites likely contribute to novel function in both of these recombinant transporters, specific amino acids at key sites are still likely context-dependent, which makes functional evolution both more difficult to predict and to engineer in this family.

Compounding this difficulty is the cryptic nature of sites that we empirically determined to influence maltotriose transport but which are unlikely to interact with the substrate (Fig. 1). These substitutions may effect subtle changes to the overall conformation of the transporter, especially where they have the potential to interact with other protein regions that are proximal in tertiary space (e.g. F468). Moreover, there is a growing appreciation that, in yeast monosaccharide sugar porters, the fine-scale environment around the substrate binding site plays a surprisingly large role in sugar recognition and specificity, both by shaping an accommodating binding pocket and through interactions between substrate-interacting and non-interacting residues within van der Waals distance (Kasahara et al. 2009; Drew et al. 2021).

In MalT434, more concrete hypotheses can be made about the molecular contributions of other sites important for novel substrate transport. Molecular docking analyses place the maltotriose ligand in close proximity to the key sites on TMH 7 and TMH 11 (Fig. S8), with several of the sugar hydroxyl groups capable of engaging in a hydrogen-bonding network with the side chains of polar amino acid residues at those sites. Of the substitutions in MalT434 that face the transport channel, all three have polar and hydrogen bond-competent side chains of small-to-medium size; in wild-type MalT4, the residues at these sites have bulkier and/or hydrophobic side chains. Similarly, at the crucial 379 site on TMH 7, the permissive serine has a much smaller side chain than the prohibitive glutamine. Either of the prohibitive residues at 379 and the other crucial site 505 might introduce steric clashes with the terminal glucopyranose moiety of maltotriose (Fig. S8c), even though they themselves are likely capable of hydrogen-bonding with the substrate. Notably, the residue at position 379 may be involved in coupling substrate binding to gating during the transition to the occluded state (Drew et al. 2021), a key determinant of substrate recognition that involves more tightly embedding the sugar molecule in its binding site within the transport channel. In wild-type MalT4, position 379 has the smaller serine residue, while sites along TMH 11 have bulkier amino acids; in wild-type MalT3, position 379 has the larger glutamine residue, but TMH 11 has smaller, hydrophilic residues. Thus, in each native maltose transporter, the steric constraint of the transport channel may be finely tuned at co-evolving sites along TMH 7 and TMH 11 to accommodate maltose with higher affinity and specificity, which occur at the expense of steric exclusion of other substrates, such as maltotriose (Fig. S8e). This model is consistent with the crucial role of amino acid side chain length in shaping substrate specificity in some monosaccharide sugar porters (Kasahara et al. 2011; Drew et al. 2021), notwithstanding that we also detected a complex interaction between size and biochemical properties at the key 505 site.

The difficulty of functional innovation in the high-specificity Agts begs the question of how the related generalist Agts are capable of transporting not only maltose and maltotriose, but a diverse range of substrates. If the generalist transporters had evolved from a more specific ancestor, as has been suggested (Pougach et al. 2014), their extant substrate range would imply multiple bouts of highly constrained functional evolution. To determine when and how this broad substrate specificity may have evolved in the generalist Agts, we reconstructed the yeast sugar porter phylogeny from 332 newly annotated, representative Saccharomycotina genomes encompassing more than 400 million years of evolution (Fig. S5). This analysis showed that, somewhat unexpectedly, the high-specificity Agts are a derived clade within the generalist-like Agts (Fig. 6a). The copy number of these putative generalist Agts encoded by yeast genomes is strongly predictive of growth on Agt1-exclusive substrates (Fig. 7), which further supports the conclusion that these proteins are likely bona fide generalists. The evolution of maltotriose transport by high-specificity Agts is thus better regarded as a reacquisition of ancestral function than the de novo evolution of a truly novel function within this protein family.

It remains subject to debate whether the general trend of protein evolution is directional: from less to more intrinsically specific (Bridgham et al. 2006; Tawfik 2010; Copley 2012; Steindel et al. 2016; Wheeler et al. 2016; Wheeler and Harms 2021). Multiple lines of evidence now suggest that this mode is dominant in genes involved in α-glucoside metabolism in yeasts. In addition to the α-glucoside transporters, both the α-glucosidases of *S. cerevisiae* and the transcriptional activators that regulate the structural metabolic genes likely evolved from promiscuous ancestral proteins that optimized subfunctions following duplication events, rendering them specific for different α-glucosides (Brown et al. 2010; Voordeckers et al. 2012; Pougach et al. 2014). The extent of intramolecular epistasis apparent in the high-specificity Agts, which may arise both from intra-protein and protein-substrate interactions, may provide an explanation for the inherent difficulty of re-evolving maltotriose transport in these proteins. Functional entrenchment by historical contingency and epistasis is well documented, and the irreversibility of evolutionary trajectories at the molecular level may be a widespread phenomenon (Ortlund et al. 2007; Bridgham et al. 2009; Soylemez and Kondrashov 2012; Harms and Thornton 2014; Bank et al. 2015; Podgornaia and Laub 2015; Shah et al. 2015; Starr and Thornton 2016; Starr et al. 2017; Starr et al. 2018; Ben-David et al. 2020; Xie et al. 2021; Park et al. 2022). Although not directly tested here, there may be inherent tradeoffs between specificity and substrate affinity in yeast Agts (Stambuk and Araujo 2001; Salema-Oom et al. 2005; Hatanaka et al. 2022), which would suggest that walking back through the accumulated mutations that led to higher specificity in the Mal31-like transporters would be likely to incur an immediate functional tradeoff and therefore fitness cost. The recurrent gene conversion events that enable maltotriose transport among members of this family may, therefore, represent the only meaningfully accessible route to bypass these deleterious intermediates, but the high degree of context-dependence for mutational effects makes the prediction or engineering of this novel function difficult (Hatanaka et al. 2022).

Might the evolution of yeast sugar porters more broadly be organized along an axis of increasing specialization and specificity? This family encompasses functionally diverse transporters with varying specificities for different mono- and di-saccharides and sugar alcohols; notably, functionally similar proteins are not monophyletic across the family (Donzella et al. 2023). Our phylogenetic analysis of these proteins places the Agts, which may retain some glucose transport capacity (Wieczorke et al. 1999), as a deeply branching sister clade to most of the broader family (Fig. S5). These results imply multiple bouts of functional specialization from a highly promiscuous ancestor, in some cases starting from partially subfunctionalized ancestral proteins, with the Agts perhaps remaining the most representative of the ancestral multifunctionality. While the extant diversity of yeast sugar porters has generally been regarded as an example of functional diversification (i.e. highly plastic gains of novel substrate affinity; (Brown et al. 2010; Hatanaka et al. 2022; Donzella et al. 2023)), the evolution of this important gene family may have followed a very different mode. In the former model, functional diversification by neofunctionalization follows duplication of ancestral transporter genes, whereas our analyses suggest that duplications in this gene family may be primarily followed by subfunctionalizing escapes from adaptive conflict (Hughes 1994; Hittinger and Carroll 2007; Des Marais and Rausher 2008), wherein transporters can gain increased specificity and affinity for a narrow substrate range at the expense of other ancestral ligands.

These two models have distinct implications for the myriad biotechnological applications predicated upon sugar consumption by yeasts, which might be targets for improvement by protein engineering. If extant transporters are indeed highly plastic and evolvable, shifting or expanding their substrate range should be relatively simple. If, on the other hand, they have undergone entrenched specialization, they may be inherently less evolvable (Bridgham et al. 2009; Starr et al. 2018; Wheeler and Harms 2021). Results here and elsewhere (Hatanaka et al. 2022) support the latter corollary. However, this model also implies that reconstructed ancestral proteins, or even generalist extant proteins from this clade, might both possess desirable properties and be inherently highly amenable to engineering, mutagenesis, or directed evolution approaches.

## METHODS

### Strains and cultivation conditions

*S. eubayanus* strains, plasmids, and oligonucleotides used in this work are listed in Tables S1 and S2. Yeasts were propagated on YPD medium (1% yeast extract, 2% peptone, 2% glucose) supplemented with 400mg/L G418 and/or 50mg/L Nourseothricin (CloNAT) as appropriate and cryopreserved in 15% glycerol at −80° for long-term storage.

Transformation of *S. eubayanus* was performed by the PEG/LiAc/carrier DNA method (Gietz and Schiestl 2007) with minor modifications (Baker and Hittinger 2019). CRISPR-mediated gene deletions and insertions were achieved by co-transformation of pXIPHOS vectors (Kuang et al. 2018) and repair templates for homologous recombination. Repair templates were purified PCR products consisting of single linear fragments, multiple linear fragments for *in vivo* assembly, or recombinant amplicons generated by overlap extension PCR, depending on the application. All repair templates were amplified using Phusion polymerase (New England Biolabs) per the manufacturer’s instructions and purified using QiaQuick or MinElute spin columns (Qiagen).

We assessed transporter function via expression from the native *MALT4* locus in yHJC207, a haploid derivative of the wild strain yHKS210 that was constructed as previously described (Crandall et al. 2023). Because the *MALT2* and *MALT4* loci are recent duplicates and almost identical at the nucleotide level, transporter variants were inserted into both loci out of necessity. Both *MALT2* and *MALT4* were simultaneously deleted using CRISPR-Cas9 and replaced with *kanMX*. Novel transporter variants, as well as *MALT434* and *S. eubayanus AGT1* positive controls, were subsequently inserted into both loci by co-transformation with a pXIPHOS vector expressing Cas9 and a gRNA targeting *kanMX* (Lee et al. 2021). Transformants were cured of plasmids, and the inserted alleles were sequenced.

### Quantitative growth measurements of *S. eubayanus* strains

Strains were streaked to single colonies on YPD plates, arrayed in 96-well plates in a randomized layout, and precultured in YPD at room temperature for 72 hours with gentle shaking. Precultures were serially diluted in minimal medium (0.5% ammonium sulfate, 0.017% Yeast Nitrogen Base) and inoculated into minimal medium containing 2% sugars in 96-well plates at a final dilution of 10^−4^. OD_600_ was measured every hour for 7 days using a SPECTROstar Omega plate reader (BMG Labtech) equipped with a microplate stacker. Raw growth data was summarized using GCAT (Bukhman et al. 2015). Area under the curve (AUC) measurements for growth on maltotriose, normalized to a common negative control within each experiment, were used as a response variable in linear models with protein identity (MalT3 or MalT4) at each domain or at key amino acid sites as categorical predictor variables. The effects of protein identity at some single regions and for many pairwise interactions could not be estimated due to singularities. We tested for evidence of epistasis by statistically comparing additive models and those with interaction terms (Li and Fay 2019). The amino acid properties compiled to test associations with transporter function included chemical composition, polarity, and volume (Grantham 1974), aromaticity (Xia and Li 1998), hydropathy (JANIN 1979; Kyte and Doolittle 1982; Hopp and Woods 1983; Eisenberg et al. 1984; Rose et al. 1985; Cornette et al. 1987; Engelman et al. 2003), and BLOSUM similarity (Henikoff and Henikoff 1992). Some matrices were compiled from Braun (Braun 2018). For dimensionality reduction, BLOSUM similarity was omitted.

### Quantitative growth measurements of Saccharomycotina yeasts

Growth on α-glucosides was measured for the strains whose genome annotations were analyzed, which were primarily the type strains for their respective species. Strain information, including taxonomic order (Groenewald et al. 2023), major clade (X.-X. Shen et al. 2018), and updated annotation mapping, can be found in Table S3. Cryopreserved strains were inoculated directly to YPD in 96-well plates and incubated for 7 days at room temperature. Some slow-growing species failed to revive during this time frame and were removed from further analysis, and we did not phenotype opportunistic pathogens, ultimately resulting in data for 287 species. Precultures were inoculated to minimal medium with 1% sugar or no added carbon source using a pinning tool, incubated for 7 days at room temperature, and re-inoculated to new plates containing the same medium. OD_600_ of the second round of growth was measured every hour using a SPECTROstar Omega plate reader (BMG Labtech) equipped with a microplate stacker. The growth experiments were performed four times independently. Raw growth data was summarized using Growthcurver (Sprouffske and Wagner 2016). Wells with poor model fits were discarded, and each curve was manually inspected to identify species with unreliable growth curves (Opulente et al. 2024). Growth on each carbon source was normalized to the average growth of the same species in medium with no added carbon to control for background growth. Caper (cran.r-project.org/web/packages/caper/index.html) was used to fit phylogenetically corrected regressions (PGLS) to growth data and square-root transformed Agt number, using the rooted ML species phylogeny (X.X. Shen et al. 2018).

### Structure prediction and analyses

Structural models for MalT434 were generated using four different software: AlphaFold2 (Jumper et al. 2021), Phyre2 (Kelley et al. 2015), I-TASSER (Yang et al. 2015), and SWISS-MODEL (A. Waterhouse et al. 2018). All gave extremely similar results across the structured region (mean and SD pairwise RMSD: 1.61±0.51Å), and AlphaFold2 models for all proteins of interest were generated and used for further analysis. Docking of maltotriose was performed using SwissDock (Grosdidier et al. 2011). Structure models and docking results were visualized in PyMol v2.5 (Schrödinger, LLC).

### Genome annotation

To improve the quality of existing gene models, publicly available genome assemblies of 332 Saccharomycotina yeast species (X.-X. Shen et al. 2018) were re-annotated de novo. For consistency, we retained the assembly and species names, although some species have since been renamed; consult MycoBank (www.mycobank.org) for the most up-to-date taxonomic information. Repetitive sequences were softmasked with RepeatMasker v4.1.2, and protein-coding genes were annotated using ab inito predictors AUGUSTUS v3.4.0 (Stanke et al. 2008) and GeneMark-EP+ v4.6.1 (Brůna et al. 2020) in BRAKER (Brůna et al. 2021), with Saccharomycetes proteins in OrthoDB v10 (Kriventseva et al. 2019) as homology evidence and using the --fungus mode. Where applicable, the longest transcript of each gene was retained. BUSCO v5.7.0 (Manni et al. 2021) was used to assess the completeness of the new and preexisting genome annotations using single-copy yeast orthologs in OrthoDB v10 (R.M. Waterhouse et al. 2018).

This approach was chosen so as to generate a useful community resource in two ways: first, to enable direct comparisons with a larger, partially overlapping dataset of yeast genomes published recently (Opulente et al. 2024), which were annotated using identical methods; and second, to facilitate future studies by significantly improving the quality of annotations for the widely-used 332-genomes dataset. Median annotation completeness was increased from 94.6% to 98.8%, while the median percentage of missing BUSCO genes decreased to 0.9% from 4.6% (both *p* < 2.2×10^−16^, two-sided *t*-tests; Fig. S10). Table S4 documents BUSCO analyses of existing and updated annotations for all genomes. The full updated annotations in protein and nucleotide FASTA, GFF3, and GTF formats will be available on figshare (confidential link will be updated to a public link prior to publication).

### Phylogenetic analyses

The amino acid translations of the newly predicted protein-coding genes were queried by BLASTp+ v2.9 (Camacho et al. 2009) using characterized *Saccharomyces cerevisiae* sugar transporters (Mal31, Agt1, Gal2, Hxt1-5, Hxt7) retrieved from SGD (Wong et al. 2023). BLAST subjects less than 400 or greater than 1000 amino acids in length were discarded to remove partial or fused annotations, based on distributions of sugar porter length in TCDB (Saier et al. 2006; Saier et al. 2021). Remaining proteins were annotated with their most similar *S. cerevisiae* homolog using a reciprocal BLASTp search against all translated ORFs in *S. cerevisiae*, which were retrieved from SGD. Protein sequences were aligned using the E-INS-i strategy of MAFFT v7.222 (Katoh et al. 2002; Katoh et al. 2005; Katoh and Standley 2013), and the alignment was trimmed with trimAL v1.4.22 (Capella-Gutiérrez et al. 2009) using the --gappyout parameter. The phylogeny was inferred using IQ-TREE v2.2.2.7 (Minh et al. 2020) with 1000 bootstraps (Hoang et al. 2018) and automatic substitution model selection (Kalyaanamoorthy et al. 2017). Due to the significant homology between MFS proteins, this dataset contained a small proportion of non-sugar porter MFS proteins, primarily belonging to the drug:proton antiporter family. These were retained in the alignment and tree inference to test the assumption of sugar porter monophyly. As expected, the sugar porters and non-sugar porter MFS proteins formed well-supported reciprocally monophyletic clades. The α-glucoside transporter phylogeny was refined by re-aligning the proteins from that clade and inferring the phylogeny as before, albeit with 10 independent runs of IQ-TREE with 10000 bootstrap replicates each and secondary branch support assessment by SH-aLRT tests. Trees were visualized and annotated in iTOL (Letunic and Bork 2021).

## ACKNOWLEDGMENTS

We are grateful to John F. Wolters for feedback on analyses and extensive curation of databases, Dana A. Opulente for advice on phenotyping, Kaitlin J. Fisher for sharing yeast strain copies that enabled high-throughput phenotyping, Xing-Xing Shen for advice on running IQ-TREE, and the Hittinger and Sato Labs for helpful discussion.

## AUTHOR CONTRIBUTIONS

J.G.C. and C.T.H. conceived and designed the study. X.Z. performed and assessed genome annotations with input from A.R. J.G.C. performed all experiments and analyses and wrote the manuscript with input from all authors. J.G.C., A.R., and C.T.H. secured funding.

## DATA AVAILABILITY

New genome annotations for Saccharomycotina species are available on figshare (confidential link will be updated to a public link prior to publication). This confidential link is provided for review purposes and will be updated to a public link prior to publication. Other data underlying this article are available in the article and in its online supplementary material.

## SUPPLEMENTAL MATERIALS

**Table S1.** *S. eubayanus* strains and plasmids used in this study.

**Table S2.** Oligonucleotides used in this study.

**Table S3.** Strain information for the 332 Saccharomycotina species. Column A (“Species name”) corresponds to Column C (“Species name”) of Table S1 from X.-X. Shen et al. 2018.

**Table S4.** Benchmark Universal Single-Copy Orthologs (BUSCO) statistics for existing and updated genome annotations of species in this study.

**Data S1.** Maximum likelihood phylogenetic trees of sugar porters and outgroup MFS proteins from Saccharomycotina genomes in Newick format.

**Data S2.** Consensus phylogenetic tree of Agt clade proteins in Newick format. Branch supports are from SH-aLRT test and ultrafast bootstrapping, respectively.

**Data S3.** Maximum likelihood phylogenetic tree of Agt clade proteins in Newick format. Branch supports are from SH-aLRT test and ultrafast bootstrapping, respectively.

