## Supplementary Figures for "Specialization restricts the evolutionary paths available to yeast sugar transporters"

**SUPPLEMENTAL FIGURES**

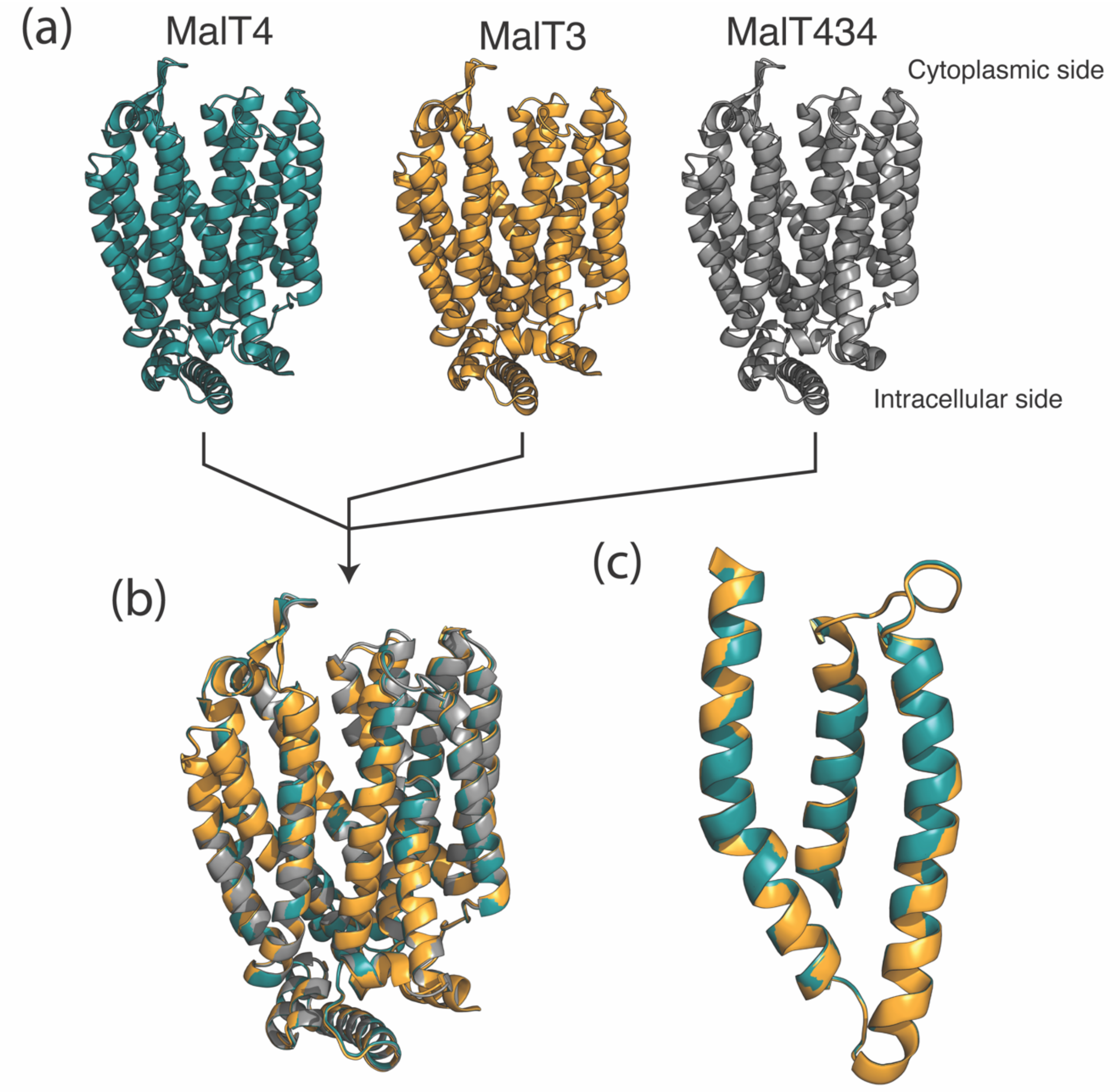


**Figure S1.** Structural similarity of wild-type and chimeric *S. eubayanus* transporters. (a) Structural models for wild-type MalT4 (teal) and MalT3 (orange), and the chimeric MalT434 (gray) are shown with intracellular N- and C-terminal domains hidden for clarity. (b) Structural overlay of MalT3, MalT4, and MalT434. The structures are superimposable with mean and SD pairwise RMSD=0.955±0.107Å. (c) Structural overlay of transmembrane helices 10, 11, and 12 from MalT3 and MalT4, corresponding to the recombinant region in MalT434 (RMSD=0.909Å).

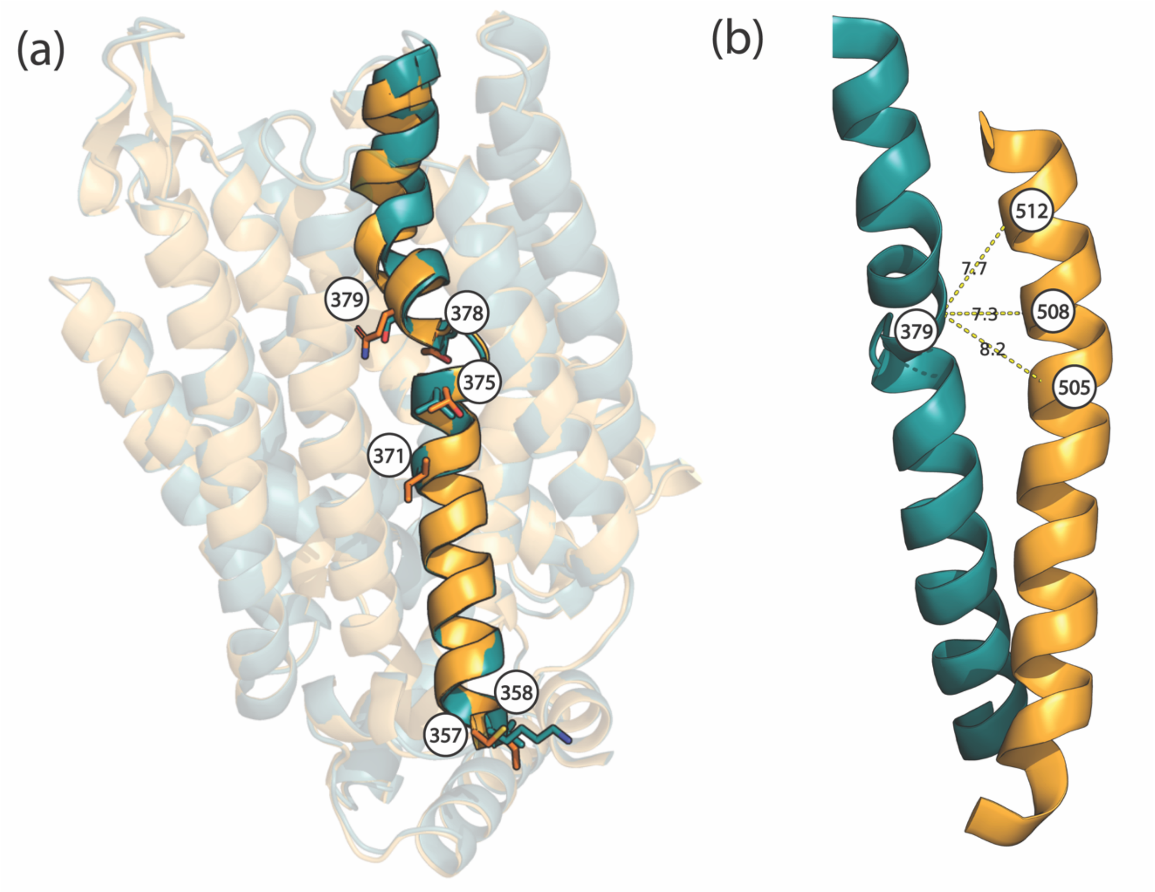


**Figure S2.** Amino acid differences between MalT3 and MalT4 along the key transmembrane helix 7. (a) The structural models of MalT4 (teal) and Malt3 (orange) are superimposed; transmembrane helix 7 is shown in full color, while all other transmembrane helices and the ICH domain are shown with transparency, and the intracellular N- and C-terminal domains are hidden. Side chains are drawn at the sites where MalT4 and MalT3 differ along TMH 7, with residue numbers labeled. (b) Detail of the proximity between position 379 on TMH 7 and sites on TMH 11. Helices are shown from the structural model of the chimeric transporter MalT434 and colored according to parental protein (TMH 7, MalT4, teal; TMH 11, MalT3, orange). Dotted lines and labels show the distance between residues 379 and 505, 508, and 512 (bottom to top, respectively), in angstroms.

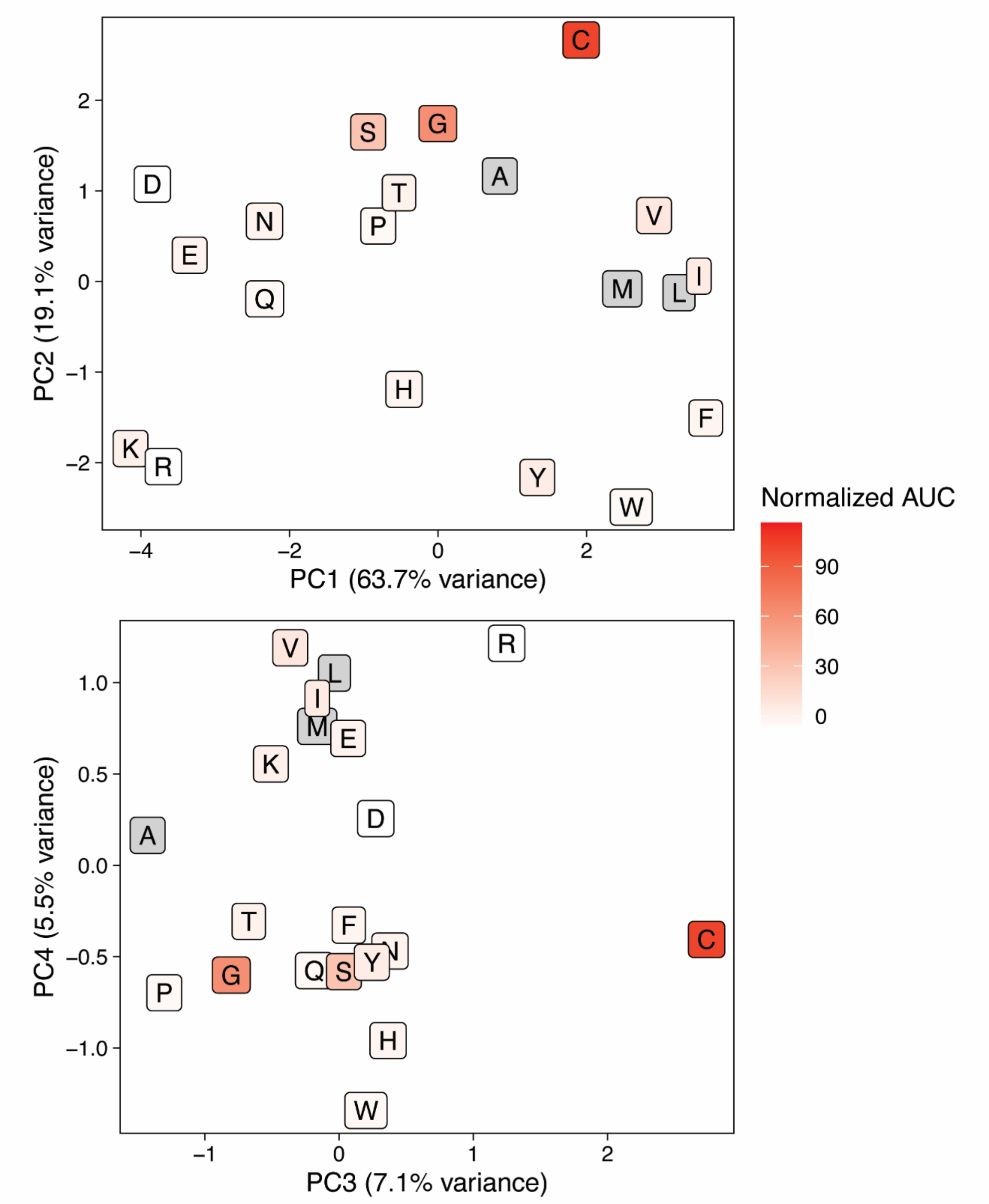


**Figure S3.** The effect of residue C505 on maltotriose transport is due to its unique properties. Scatterplots show amino acids in principal component space based on dimensionality reduction of side chain properties. Colors show growth on maltotriose (AUC, area under the curve) of strains expressing MalT4 with the given amino acid at position 505, in the context of all other necessary mutations for maltotriose transport, as shown in Fig. 5. Amino acids in gray were not measured.

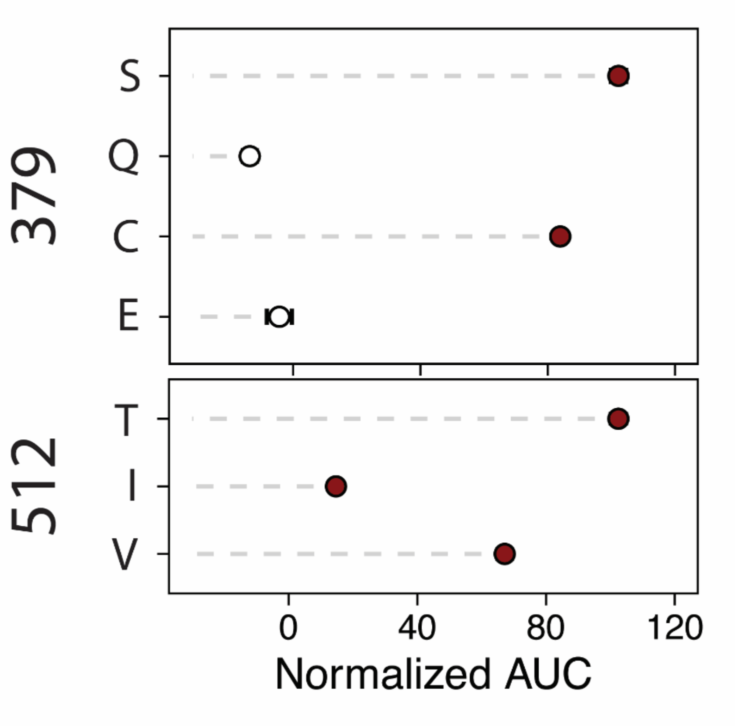


**Figure S4.** Further evidence for nuanced and context-specific biochemical requirements at key sites. Points and bars show mean +/- SEM of normalized growth on maltotriose (AUC, area under the curve) of strains expressing MalT4 variants which each contain all other necessary mutations for maltotriose transport. The amino acid identity at position 379 (top panel) or 512 (bottom panel) is shown on the x-axis. Filled circles denote growth significantly greater than the negative control ($p < 0.01$, Mann-Whitney *U* test with Benjamini-Hochberg correction).

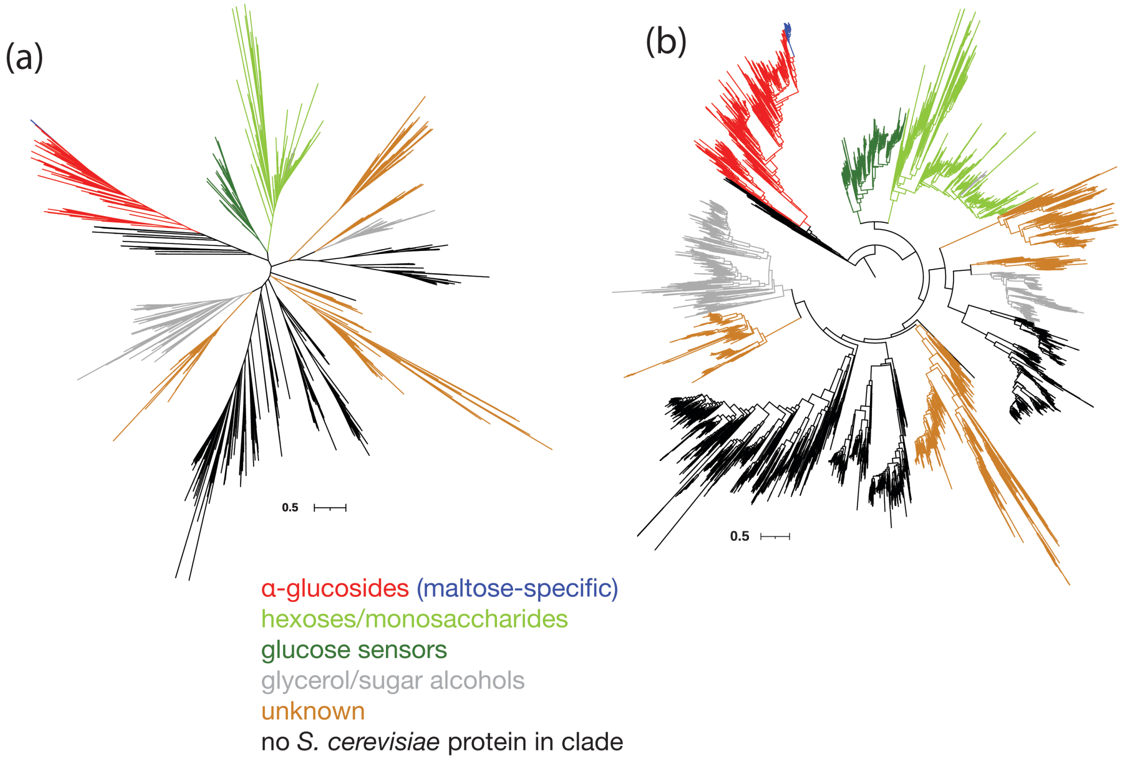


**Figure S5.** Unrooted (a) and midpoint-rooted (b) maximum-likelihood phylogeny of 8,403 sugar porters from 332 budding yeast genomes. An outgroup clade containing non-sugar porter Major Facilitator Superfamily proteins is hidden for clarity; rooting the tree to this outgroup recovers the same topology. Major clades containing at least one *S. cerevisiae* protein are colored according to the substrate(s) of the *S. cerevisiae* homolog: α-glucosides (red and blue), hexoses/monosaccharides (light green), glucose sensors (dark green), glycerol/sugar alcohols (gray), and undetermined (orange). The high-specificity maltose transporter clade, which contains the *Saccharomyces*-specific Mph2/3 proteins, is shown in blue. The Newick-formatted tree is available in Data S1.

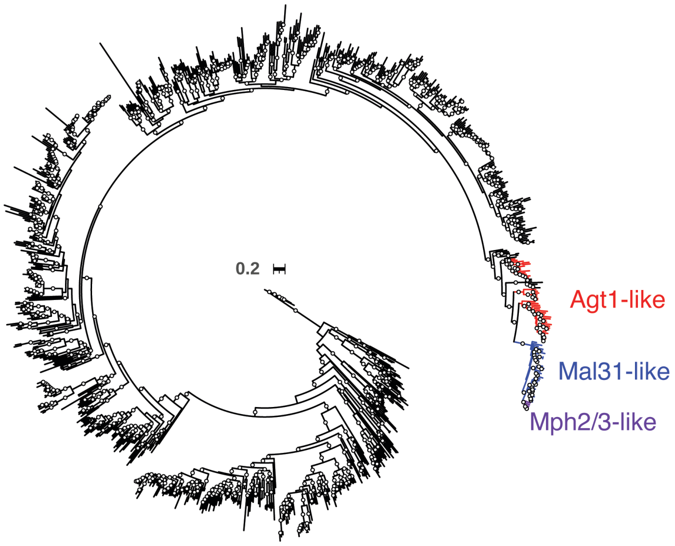


**Figure S6.** Maximum-likelihood phylogeny of the clade containing α-glucoside transporters, rooted to a divergent outgroup. Agt1- and Mal31-like transporters from Saccharomycetales are colored in red and blue, respectively, with the *Saccharomyces*-specific Mph2/3 clade in purple. Circles denote branches with >90% bootstrap support. The Newick-formatted tree is available in Data S3.

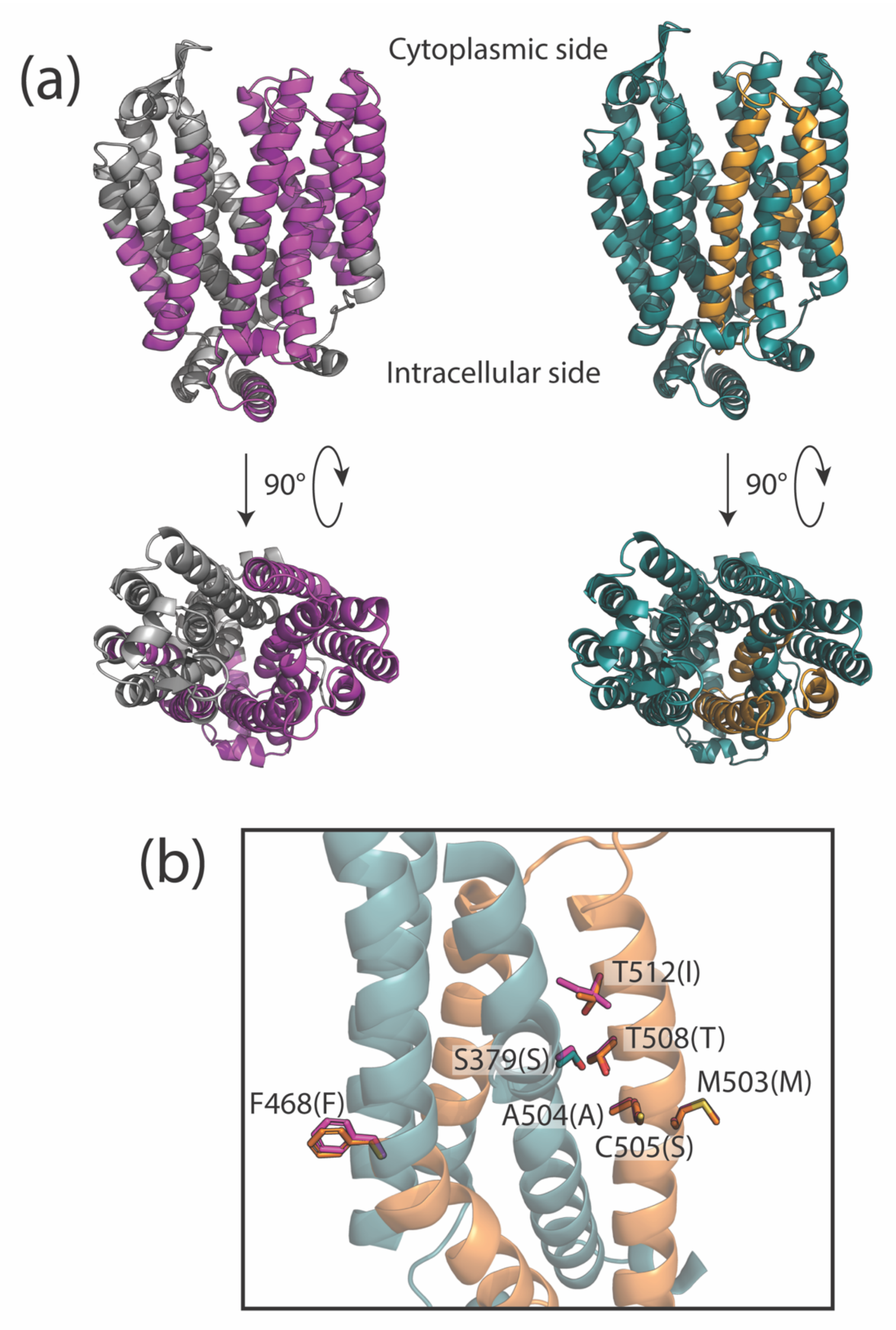


**Figure S7.** Similarity of chimeric maltotriose transporters. (a) Structural models for Mty1 (left, gray and pink) MalT434 (right, teal and orange) and are shown from side-on (top row) and top-down (bottom row) views. Intracellular N- and C-terminal domains are hidden for clarity. On each structure the contrasting colors denote regions from different parental proteins (i.e. chimeric breakpoints). The two models are superimposable with RMSD=0.898Å. (b) Detail of amino acid similarity between MalT434 and Mty1. The structural model of MalT434 is shown, with a focus on the C-terminal helical bundle as viewed from the transport channel. N-terminal helices are hidden for clarity. Side chains are drawn for residues in MalT434 that were determined to have an effect on maltotriose transport in molecular experiments, colored according to their parental protein identity (MalT4, teal; MalT3, orange). Side chains are drawn in pink for the corresponding residues in the aligned structural model of Mty1, which is hidden for clarity; the amino acid identity for Mty1 is in parentheses.

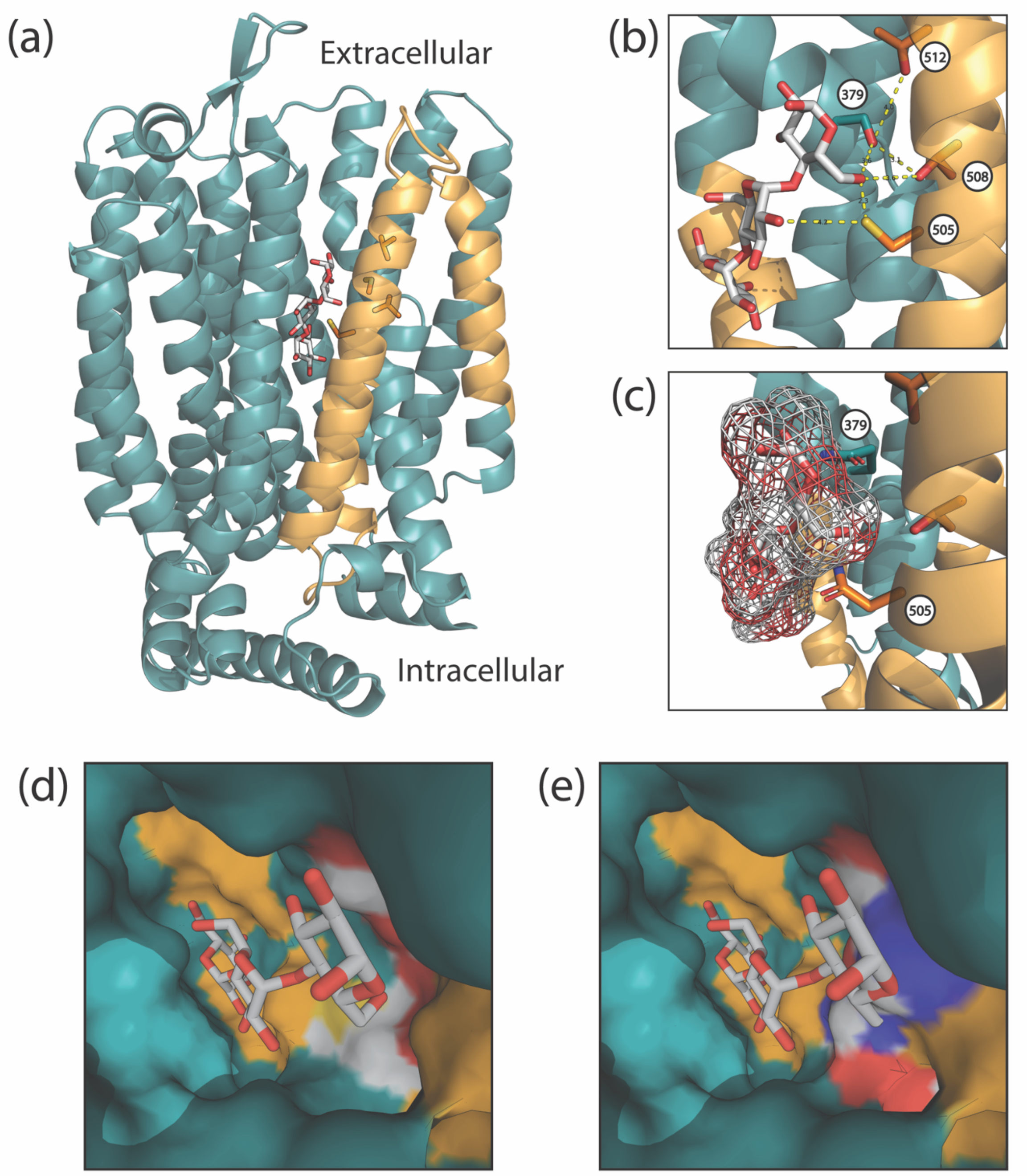


**Figure S8**. (a) A side view of the structural model of MalT434 is shown with side chains drawn at residues 379, 505, 508, and 512. Docked maltotriose is drawn as sticks and colored by element. (b) Details of proximities and potential interactions between key residues and maltotriose. The structural model with docked ligand is shown as in (a), with obscuring helices hidden. Dotted lines show distances in angstroms between maltotriose and key residues with a functional impact on maltotriose transport that could engage in a hydrogen-bonding network. (c) Residues Q379 and N505 could inhibit sugar transport by occlusion. Maltotriose docked to the structural model of MalT434 is shown as in (b) with a mesh surface drawn for the sugar. Side chains are drawn for residues S379 and N505, both of which could inhibit the accommodation of the large maltotriose molecule. (d-e) Docked maltotriose in the space-filling model of MalT434. The surface is colored by MalT3/MalT4 identity, except for at residues 379, 505, 508, and 512, which are colored by element. In (e), the residues at these positions have been reverted to their MalT3 (379) or MalT4 (505, 508, 512) identity, which substantially reduces the volume of the substrate pocket and introduces steric clashes with the sugar.

Fig. S9

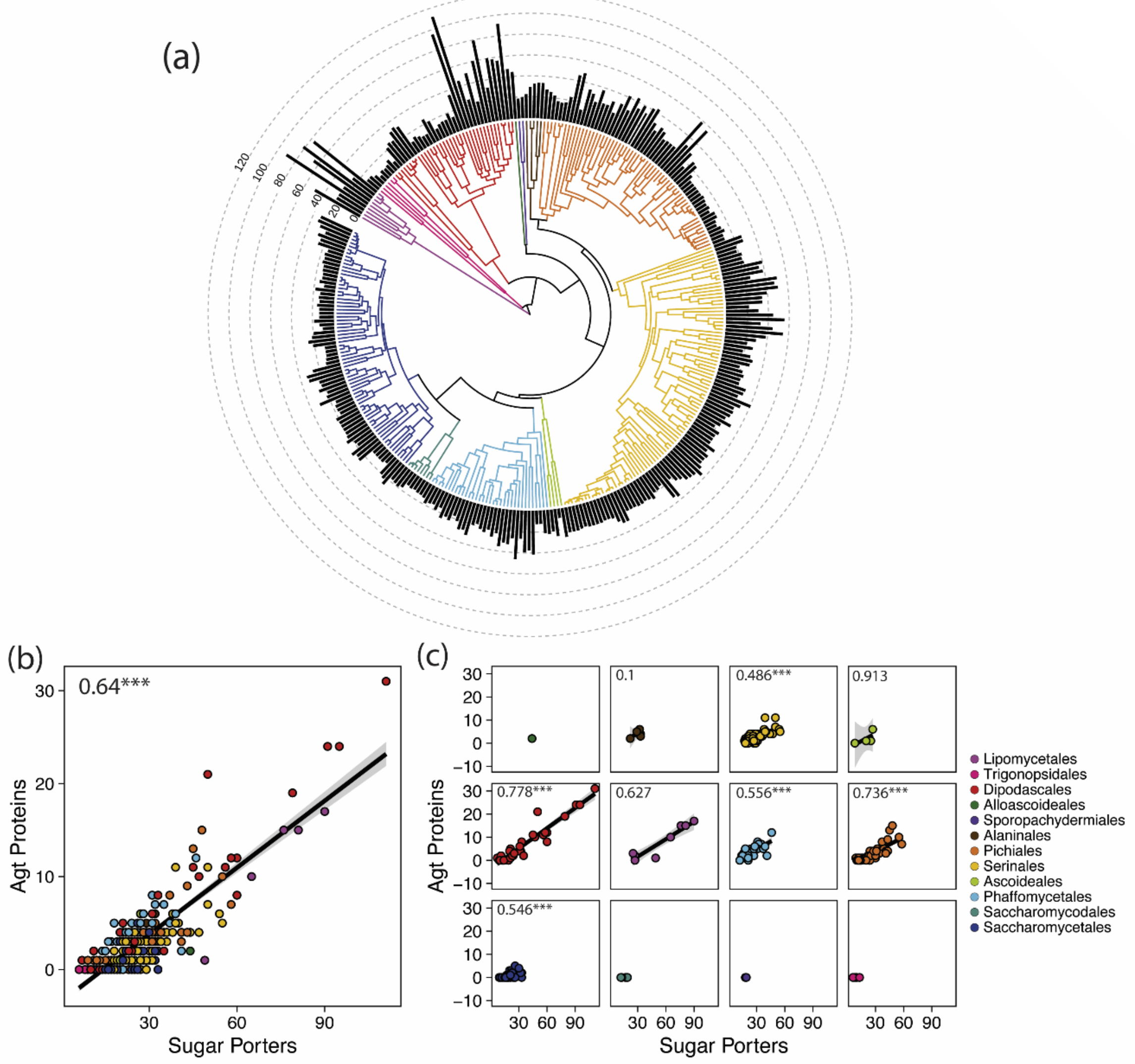


**Figure S9.** Expanded sets of Agt proteins are generally representative of genome-wide sugar porter complement. (a) Time-calibrated Saccharomycotina species phylogeny, as in Fig. 7, with branches colored by order. The bar chart shows the total number of sugar porters in each genome. (b) Scatterplot of all sugar porters versus Agt proteins encoded in each genome. Each species is a point, colored by taxonomic order. Inset text gives

Kendall's T ($p < 2.2x10^{-16}$). (c) Scatterplots as in (b), split by taxonomic order. Inset text gives within-order Kendall's T (***$p < 1.3x10^{-5}$). Correlation coefficients were not calculated for orders with too few species in our dataset. Lines and shaded regions in (b) and (c) show simple linear regressions and 95% confidence intervals of untransformed data.

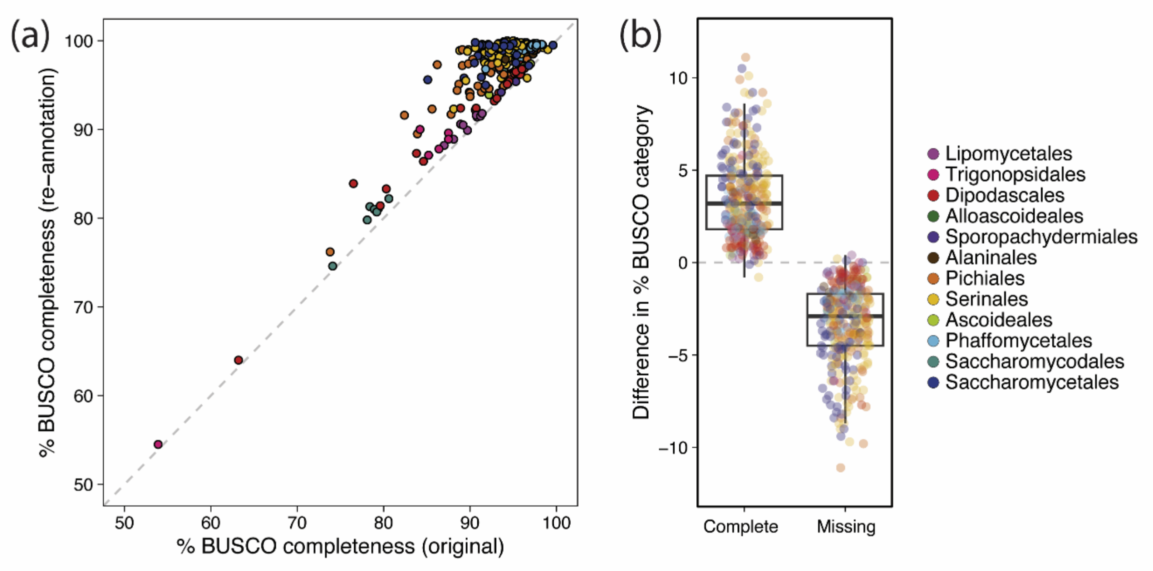


**Figure S10**. Improvement in annotation quality for 332 Saccharomycotina genomes. (a) Scatterplot of annotation completeness (% of Benchmark Universal Single-Copy Orthologs/BUSCO present in single, complete copy in annotation) for the 332 Saccharomycotina genomes used in this study (Shen et al. 2018). Completeness for previous annotations is on the x-axis, and completeness for new annotations from the current work are plotted against the y-axis. The dotted gray line shows the 1:1 null expectation of no improvement. (b) Difference in the percentage of complete and missing BUSCO for annotations from the current study compared to previous annotations. Boxplots show median and interquartile ranges; individual genomes are plotted as points. Points in both panels are colored by taxonomic order.
